# AvrSr27 is a zinc-bound effector with a modular structure important for immune recognition

**DOI:** 10.1101/2023.11.21.567997

**Authors:** Megan A. Outram, Jian Chen, Sean Broderick, Zhao Li, Shouvik Aditya, Nuren Tasneem, Taj Arndell, Cheryl Blundell, Daniel J. Ericsson, Melania Figueroa, Jana Sperschneider, Peter N. Dodds, Simon J. Williams

**Affiliations:** Plant Sciences Division, Research School of Biology. The Australian National University, Canberra, Australia; Commonwealth Scientific and Industrial Research Organisation, Agriculture and Food, Canberra, ACT, Australia; Australian Synchrotron, Macromolecular Crystallography, Clayton, Vic., 3168 Australia

**Author notes:** These authors contributed equally to this work (Megan A. Outram, Jian Chen). Corresponding authors: Peter Dodds –; Simon Williams.

**Keywords:** AvrSr27, effector recognition, metal-binding, plant immunity, Sr27, stem rust, zinc

## Abstract

Stem rust, caused by the fungal pathogen *Puccinia graminis f. sp.tritici* (*Pgt*) is a major threat for wheat production and global food security. Central to the success of *Pgt* is the secretion of proteinaceous effectors that promote infection and colonisation, while immunity in wheat is driven by receptor-mediated recognition of these effectors resulting in pathogen avirulence. Here, we report the crystal structure of the cysteine-rich effector AvrSr27, the third experimentally derived structure of a *Pgt* effector. The AvrSr27 structure reveals a novel β-strand rich modular fold consisting of two structurally similar domains and confirms the poor prediction we obtained from the AlphaFold2-derived model. The highly prevalent cysteine residues within the protein facilitate the co-ordination of 4 zinc molecules. Utilising the structure, we show that the N-terminal domain of AvrSr27 is sufficient for immune recognition and interaction by Sr27. The 7-cys motif sequence in each AvrSr27 domain, which facilitates zinc binding, was also found in two haustorially-expressed, structurally homologous *Pgt* proteins. Remarkably, despite relatively low sequence identity, we show that these proteins can associate with Sr27 and trigger cell death in heterologous systems and wheat protoplasts, albeit weaker than AvrSr27. Collectively, our findings have important implications for the field embarking on bespoke engineering of immunity receptors as solutions to plant disease.

## Introduction

Central to the success of adapted phytopathogens is the secretion of proteinaceous effectors that modulate their host’s physiology to promote infection and colonisation. Immunity to pathogens in plants is largely driven by receptor-mediated recognition of effectors, either intra- or extracellularly [1]. Nucleotide-binding leucine-rich repeat receptors (NLRs) are predominantly responsible for intracellular immunity, triggering defence responses following effector recognition that leads to programmed cell death to limit the spread of infection [2]. Wheat steam rust, caused by the fungal pathogen *Puccinia graminis f sp. tritici* (*Pgt*), is a major threat for wheat production and global food security [3]. The emergence of highly virulent *Pgt* strains, such as the Ug99 lineage first identified in Uganda in 1998 [4, 5] has prompted a global effort to identify effective immunity receptors as genetic resistance is currently the most economical and sustainable approach for rust control. Optimum deployment of immunity receptor genes in the field depends on knowledge of the corresponding pathogen avirulence (Avr) effector proteins and their genetic diversity. To date, five *Pgt* Avr effectors, AvrSr50, AvrSr35, AvrSr22, AvrSr13, and AvrSr27, have been identified and recognition by the corresponding wheat NLRs, Sr50, Sr35, Sr22, Sr13, and Sr27, has been demonstrated [6-9].

The interaction between effectors and NLRs is a key driver for evolution and has resulted in highly sequence diverse effector proteins that typically lack conserved functional motifs or similarity to proteins of known function [10]. Structural biology and protein biochemistry approaches have provided valuable insights into the molecular functions of effectors and to elucidate the molecular mechanisms underpinning recognition and receptor activation [11]. In the *Pgt*-wheat pathosystem, structural elucidation of AvrSr50 has allowed mapping of polymorphisms controlling recognition by Sr50 to a single surface exposed residue on the Avr effector protein [12], while a phylogenetics and structural modelling approach has enabled identification of the recognition surface on the Sr50 LRR domain [13]. Recent resolution of the AvrSr35/Sr35 resistosome structure by two independent groups has revealed specific binding of the effector to the lateral ascending surface of the Sr35 LRR and subsequent activation of the NLR through a steric clash mechanism [14, 15].

At present, no structure of AvrSr27 has been reported. Three alelles of AvrSr27 have been identified (AvrSr27-1, AvrSr27-2, avrSr27-3), all of which can be recognised by Sr27, although the avrSr27-3 variant shows lower expression and does not confer avirulence in *Pgt* [7]. These AvrSr27 proteins physically associate with Sr27 *in planta* [16], suggesting recognition by direct interaction. The AvrSr27 sequence shares no sequence similarity to other proteins, but unlike AvrSr35 and AvrSr50 shows a high cysteine content (14 or 15 cysteine residues, ∼12% of the mature protein). Cysteine-rich apoplastic effectors usually contain multiple disulfide bonds but in two cysteine-rich cytoplasmic effectors with known structures, the cysteines are involved in zinc ion coordination: AvrP from the flax-rust pathogen *Melampsora lini* [17] and AvrPii from the rice blast pathogen *Magnaportha oryzae* [18]. Here, we show using an *E. coli*-based expression and purification workflow [19] that AvrSr27 also binds to zinc ions via its cysteine residues. We determined the crystal structure of AvrSr27 using a zinc-phasing approach and show it has a novel modular fold consisting of a duplicated domain, with the N-terminal domain sufficient for Sr27 recognition and interaction. Using a structure-informed metal-binding motif search we identified two additional haustorially-expressed *Pgt* proteins, predicted to adopt the same fold as AvrSr27, that also interact with and activate the Sr27 receptor, although more weakly than AvrSr27.

## Materials and methods

### Vector construction

For protein expression and purification, the cDNA of three *AvrSr27* variants, *AvrSr27-1*, *AvrSr27-2* and *avrSr27-3* [7], were PCR-amplified, excluding their signal peptide as identified by SignalP-5.0 [20], and cloned into a modified pOPIN expression vector using Golden Gate cloning (Table S1) [21]. Primers for amplification consisted of a gene-specific sequence flanked by overhangs compatible for Golden Gate cloning (Table S2) and digestion/ligation reactions and cycling were carried out as previously described [22]. The resulting constructs contained an N-terminal His6-tag to facilitate protein purification by affinity chromatography followed by a 3C protease cleavage site.

Sr27 and the three AvrSr27 variants constructs used for transient expression in *Nicotiana benthamiana* are as described in Upadhyaya et al. 2021 [7]. Pgt21-027343 and Pgt21-028479, excluding their signal peptide as identified by SignalP-5.0 [20], were cloned into pBIN19-YFP-GWY vector, using Gateway cloning LR-clonase reaction. Truncated fragments of AvrSr27 variants and structural homologues were PCR-amplified, excluding the signal peptide (28 amino acids) or first 34 amino acids, and cloned into pDONR207 and then into the pBIN19-YFP-GWY expression vector as described above (Table S2). For plant two-hybrid assays, the cDNA of Sr27, AvrSr27 variants and their truncated fragments were cloned into pAM-35S-GAL4BD-GWY and pAM-35S-VP16-GWY vectors using Gateway cloning [16], respectively (Table S2). Protoplast expression constructs in the pTA22 vector backbone used in this study are described in Arndell et al 2023 [9]. All vectors generated were sequence verified and a full list of constructs used in this study can be found in Table S2.

### Heterologous expression and protein purification of the three AvrSr27 variants in *E. coli*

For heterologous expression, our previously developed workflow [19] was utilised for initial expression trials for AvrSr27-1. AvrSr27-1 was transformed into either SHuffle T7 Express lysY Competent *E. coli* (C3030J) or BL21 T7 Express lysY competent *E. coli* (NEB, C3010I). Bacterial cultures were grown in Terrific Broth (TB) media at 37°C with shaking at 220 RPM until OD_600_ was 0.6-0.8, and then induced with 200 µM isopropyl-1-thio-β-d-galactopyranoside (IPTG). After induction, cultures were incubated for a further 16 h at 20°C and then harvested by centrifugation. Cells were resuspended in a lysis buffer containing 50 mM HEPES pH 7.5, 500 mM NaCl, 10% glycerol, 1 mM phenylmethanesulfonyl fluoride (PMSF) with 1 µg/g of cell pellet DNase I (Sigma). For the BL21 grown cells, 50 µM ZnCl_2_ and 1 mM dithiothreitol (DTT) was supplemented into the lysis buffer. The following steps were all carried out independently between the two cultures, and 1 mM DTT was added to all buffers for the BL21 preparation. Cells were lysed on ice by sonication (40% amplitude, 10 s on, 10 s off), followed by clarification of the lysate by centrifugation at 18,000 xg for 35 min at 4°C. The clarified lysate was applied to a 5 mL HisTrap FF excel column (GE Healthcare) and then washed with 20 column volumes (c/v) of buffer consisting of 50 mM HEPES pH 7.5, 500 mM NaCl, and 30 mM imidazole. The proteins were eluted using a linear gradient from 30 mM to 250 mM imidazole over 10 c/v. The fractions were visualised by Coomassie-stained SDS-PAGE, and those containing the protein of interest were pooled and buffer exchanged in 50 mM HEPES pH 7.5 with 500 mM NaCl, using a 3 kDa MWCO Amicon centrifugal concentrator (Merck) and the N-terminal His6-tag cleaved with Human Rhinovirus 3C protease at 4°C overnight. Complete cleavage by 3C was visualised by Coomassie-stained SDS-PAGE and the cleaved sample was purified further by size exclusion chromatography (SEC) on a Superdex 75 16/600 column (GE Healthcare) pre-equilibrated with 20 mM HEPES pH 7.5, and 150 mM NaCl. Protein concentration was determined by measuring the absorbance at 280 nm using an extinction coefficient as calculated by the Expasy ProtParam tool; http://ca.expasy.org/cgi-bin/protparam).

For crystallisation, AvrSr27-1, -2 and -3 were expressed in BL21 T7 Express lysY competent *E. coli* (NEB, C3010I), and purified as described above, with the exception that 50 µM ZnCl_2_ was added to the TB media during the expression step to ensure full loading of the protein with zinc ions. EDTA (1 mM final concentration) was added to the sample immediately prior to loading onto SEC and the running buffer for SEC was 20 mM HEPES pH 7.5, 150 mM NaCl, 1 mM DTT, and 50 µM ZnCl_2_.

### Intact Mass Spectrometry

The monoisotopic mass of avrSr27-3 in a native, alkylated, and EDTA/alkylated state was measured on an Orbitrap Fusion Tribrid Mass Spectrometer (ThermoFisher Scientific) coupled to Dionex Ultimate 3000 UHPLC, following liquid chromatography and heated electrospray ionisation with the results analysed using Freestyle software. Liquid chromatography was performed using a 15-minute acetonitrile gradient on an Agilent Zorbax 300SB-C3 Rapid resolution 4.6 x 50 mm 3.5 micron analytical column at a flow rate of 0.5 μL/min. For the native sample, AvrSr27 at a molar concentration of 10 μM (diluted in formic acid (FA; final concentration of 0.1% v/v)) was used. For the alkylated sample, ∼100-500 μM of protein was incubated with 20 mM iodoacetamide (IAA) for 30 mins at room temperature in the dark, followed by a 1:50 dilution for BL21-AvrSr27 and a 1:20 dilution for SHuffle-AvrSr27 with 0.1% v/v FA (∼10 μM final concentration). For the EDTA treated sample, 500 μM of avrSr27-3 was incubated in 1 mM of EDTA for 30 mins at room temperature and treated with IAA before dilution in FA as described above. For each sample, 5-10 μL (total protein amount ∼0.7 μg) was loaded for measurement. Theoretical monoisotopic masses were calculated using the Expasy PeptideMass program (https://web.expasy.org/peptide_mass/).

### 4-(2-pyridylazo) resorcinol (PAR) assay

PAR assays were carried out as described previously [19]. Briefly, the PAR solution was prepared in MilliQ water. Proteins were boiled with 1% SDS for 10 min for denaturation and metal ion release. The denatured proteins were mixed with the PAR solution to a final concentration of 50 μM PAR and 15 μM of protein. For blank measurements with buffer, the protein solution was substituted with an equal buffer volume. The absorbance of the samples was measured at wavelengths between 250 and 700 nm using a Spectromax Quickdrop (Molecular Devices).

### Metal ion characterisation and quantification via ICP-MS

Metal ions associated with purified proteins were characterised following their expression in BL21 in TB media supplemented with a 1000X trace elements solution (50 mM FeCl_3_, 20 mM CaCl_2_, 10 mM MnCl_2_, 10 mM ZnSO_4_, 2 mM CoCl_2_, 2 mM CuCl_2_, and 2 mM NiCI_2_) [23] using Inductively Coupled Plasma – Mass Spectrometry (ICP-MS). To prepare samples for ICP-MS analysis, proteins were purified (as detailed above) and dialysed extensively against one litre of buffer containing 20 mM HEPES pH 7.5, 150 mM NaCl. Subsequently, 5 mM of dialysed protein was incubated in 2% HNO3 (analytical grade) overnight at room temperature followed by centrifugation at 16,000 xg for 10 min. Resulting supernatants were analysed on a ThermoFisher iCap RQ Quadrupole – ICPMS in KEDS (Kinetic Energy Dispersion Sensitive) mode using helium as a collision gas at 1550 W. Argon was used to generate plasma and nebulise the samples. Agilent Intelliquant 68 multi-element standard No.1 (IQ-1, #5190-9422) at 100 mg/mL in 5% HNO3 was used to prepare standards for the calibration curve. The software Qtegra was used to process the data. Limit of detection was calculated from the standard deviation across instrument blanks in triplicates. Each experiment was carried out in triplicate.

### Microscale Thermophoresis (MST)

MST experiments were performed on a Monolith.NT115 instrument (NanoTemper Technologies, Munich, Germany) at 25 °C. AvrSr27-1 was labelled with Alexa Flour 647 succinimidyl ester (Thermo Fisher Scientific) and used at a final concentration of 100 nM. In brief, 20 μM of the protein solution was incubated with 2-fold molar excess of the fluorophore for 2 hours in the dark at room temperature, and the free dye was subsequently removed using a PD-10 desalting column (Cytiva). Stock solutions of the metal ions were serially diluted in buffer (20 mM HEPES pH 7.5, 150 mM NaCl), mixed 1:1 with the labelled protein and loaded into standard capillaries (NanoTemper Technologies). MST measurements were recorded using 20 % LED power and 20 to 80 % MST power and analysed using MO. Affinity Analysis software 2.2.7 (NanoTemper Technologies).

### Crystallisation, data collection and structure determination

Initial screening to determine crystallisation conditions for three allelic variants; AvrSr27-1, AvrSr27-2 and avrSr27-3 was performed in MRC2 96-well plates at 20°C using the sitting-drop vapour-diffusion method and commercially available sparse matrix screens. For screening, 300 nL drops, which consisted of 150 nL protein solution and 150 nL of reservoir solution, were prepared using a NT8®-Drop Setter robot (Formulatrix, USA). The drops were monitored and imaged using the Rock Imager system (Formulatrix, USA). No crystals were observed for AvrSr27-2 and avrSr27-3, but crystals of AvrSr27-1 (at a concentration of 15.6 mg/mL) were observed in many conditions. One of these conditions (0.2 M sodium acetate, 0.1 M Tris pH 8.5, and 30% w/v PEG4000; A9, SG1) was chosen for further optimisation in a 24-well hanging-drop vapour diffusion plate format, with 1 µL protein solution equilibrated against 1 µL of reservoir solution. Single crystals were obtained by streak seeding using a seed stock in 0.2 M sodium acetate, 0.1 M Tris pH 7.5, 30% w/v PEG4000, with 10 to 20% w/v glycerol.

SAD (single-wavelength anomalous diffraction) was used to determine the structure factor phases, taking advantage of the bound Zn ions in AvrSr27-1. A dataset was collected to obtain the anomalous signal at the high-energy remote wavelength (9.83 keV, 1.26 Å) on the MX1 beamline at the Australian Synchrotron [24] (Table S3). The dataset was scaled using Aimless [25] in the CCP4 suite [26]. For SAD phasing, the CRANK2 pipeline was used [27] in the CCP4 suite. The model was then refined using phenix.refine in the PHENIX package [28], and iterative model building between refinement rounds was carried out in Coot [29]. Structure validation was carried out using the MolProbity online server [30]. The coordinates and structure factors have been deposited in the Protein Data Bank with the accession number 8V1J.

Structure based comparisons to proteins of experimentally determined structures were carried out using the Dali online web server [31]. A search for the motif CX_13_CX_2_CX_11_CX_2_CX_9_CX_4_C in the *Pgt* secretome [32] was conducted using seqkit [33].

### Structural predictions

Structure predictions were conducted with AlphaFold2 v2.3.0 [34] using the full databases for multiple sequence alignment (MSA) construction. All templates downloaded on July 20, 2021 were allowed for structural modelling. For each of the proteins, we produced five models and selected the best model (ranked_0.pdb) for visualisation purposes. These models are included as a supporting file.

### Transient expression in Nicotiana benthamiana and N. tabacum

Four-week-old *N. benthamiana* and *N. tabacum* plants growing at 24L with 16 hours light were used for transient expression assays. *Agrobacterium* strains were cultured overnight at 28°C in LB media with required antibiotic selections. The cells were harvested and resuspended in Agro-infiltration buffer (10LmM MES pHL5.6, 10LmM MgCl_2_, 150 μM acetosyringone) and incubated at room temperature for 2 hours. For HR assays, the Sr27-containing *Agrobacterium* were infiltrated at a lower optical density (OD_600_) of 0.15 in *N. benthamiana* and 0.4 in *N. tabacum*, and the AvrSr27-containing *Agrobacterium* were infiltrated at OD_600_ = 0.5. Leaves were photographed 3 days after infiltration and were scored visually (on a scale from 0 to 4) to assess cell death. For plant two-hybrid assays, the GAL4BD-Sr27 containing *Agrobacterium* were infiltrated at OD_600_ = 0.8, and the VP16-AvrSr27 containing *Agrobacterium* were infiltrated at OD_600_ = 1.0. Leaves were photographed 5 days after infiltration and then treated by immersing in 100% ethanol until the leaves become decolorised.

Betalain was extracted by immersing three leaf disks (0.8 cm diameter) from three independent leaves into 0.5 ml water. The absorbance of betalain was measured at 538 nm using a spectrophotometer (Biochrom WPA).

### Western blot analysis for protein expression confirmation

For immunoblot analysis, total proteins from infiltrated leaves were extracted using Laemmli buffer and separated by SDS-PAGE and transferred to nitrocellulose membrane. For the proteins fused to a YFP tag, membranes were blocked with 5% skim milk and then incubated in the primary anti-GFP mouse monoclonal antibody (Roche, 11814460001), followed by goat anti-mouse HRP conjugate (BioRad, 1705047). For the plant two-hybrid samples, membranes were incubated in the primary anti-GAL4BD (Sigma, G3042) and anti-VP16 (Sigma, V4388) antibodies followed by anti-rabbit HRP conjugate (Promega, W4018). Protein loading was visualised by staining membranes with Ponceau S and protein signals were developed using the SuperSignal West Femto chemiluminescence kit (Pierce).

### Protoplast isolation, transformation, and flow cytometry

Wheat (*Triticum aestivum* cv. Fielder) seeds were planted in 13 cm pots (12 seeds per pot) containing Martins Seed Raising and Cutting Mix supplemented with 3 g/L osmocote. Seedlings were grown in a growth cabinet at 24°C on a cycle of 12 hours light (∼100 µmol m^-2^ s^-1^) and 12 h dark, for 7– 8Ldays. Protoplast isolation was carried out as described previously [35], except that released protoplasts were filtered through a 40 μm nylon cell strainer and final suspension was in 6 mL MMG solution (4 mM MES-KOH (pHL5.7), 0.4 M mannitol, 15LmM MgCl_2_). The protoplast concentration was determined by cell counting on a hemocytometer, and then adjusted to 2.5L×L10^5^ cells/mL using MMG solution. Transformations were performed in triplicate for all treatments and controls, as described in [9]. Three pmol of each vector (YFP reporter, Avr, and Sr27 or empty vector) was mixed with 200 μL of protoplast suspension (∼50,000 cells). Flow cytometry of transformed cells after 24-hours was performed as previously described [9].

## Results

### AvrSr27 is a metal-bound effector with a preference for binding zinc ions

With the advent of new deep learning tools for structure prediction, we first searched the AlphaFold2 (AF2) database [34, 36] to determine whether a confident model of AvrSr27 existed and found that models for the three AvrSr27 variants were substantially different, and all had low confidence scores (Fig. S1A). We also performed predictions using AF2v2.3.0 and again found substantial differences between models for the three variants, although the confidence in the models overall was higher (Fig. S1B). We therefore sought to experimentally determine the structure of the AvrSr27 proteins. Given the high cysteine-content, we followed a workflow for cytoplasmic expression of small cysteine-rich effectors in *E. coli* [19], by expressing avrSr27-3 with an N-terminal cleavable 6xHis-tag in the *E. coli* strains SHuffle® (oxidising environment) [37] or BL21 (reducing environment). The protein was purified to homogeneity using nickel affinity and size exclusion chromatography (SEC). A comparison of the avrSr27-3 peaks from SEC, and via SDS-PAGE (equal volume loaded) indicated that the overall protein yield from BL21 was ∼10X greater than from SHuffle® (Fig. 1A). This indicates that protein production is favoured under reducing conditions and suggests the cysteines are likely reduced. The monoisotopic mass for avrSr27-3 produced from either BL21 or SHuffle®, measured via intact mass spectrometry (MS), corresponded to the expected theoretical molecular mass (13518 Da) associated with the reduced form of the protein (Fig. 1B, Fig. S2A). To further confirm this observation, the BL21 produced avrSr27-3 was alkylated with iodoacetamine (IAA), which results in a mass increase of 57 Da per free thiol/thiolate group. The measured mass showed a range of peaks each 57 Da apart, suggesting that incomplete alkylation had occurred (Fig. S2B). With the possibility that bound metal ions may be hindering alkylation, we performed alkylation after pre-treatment with the metal chelating agent EDTA and observed a single peak with a monoisotopic mass of 14316 Da, consistent with complete alkylation of all the cysteine residues (Fig 1B, S2C).

**Figure 1:**
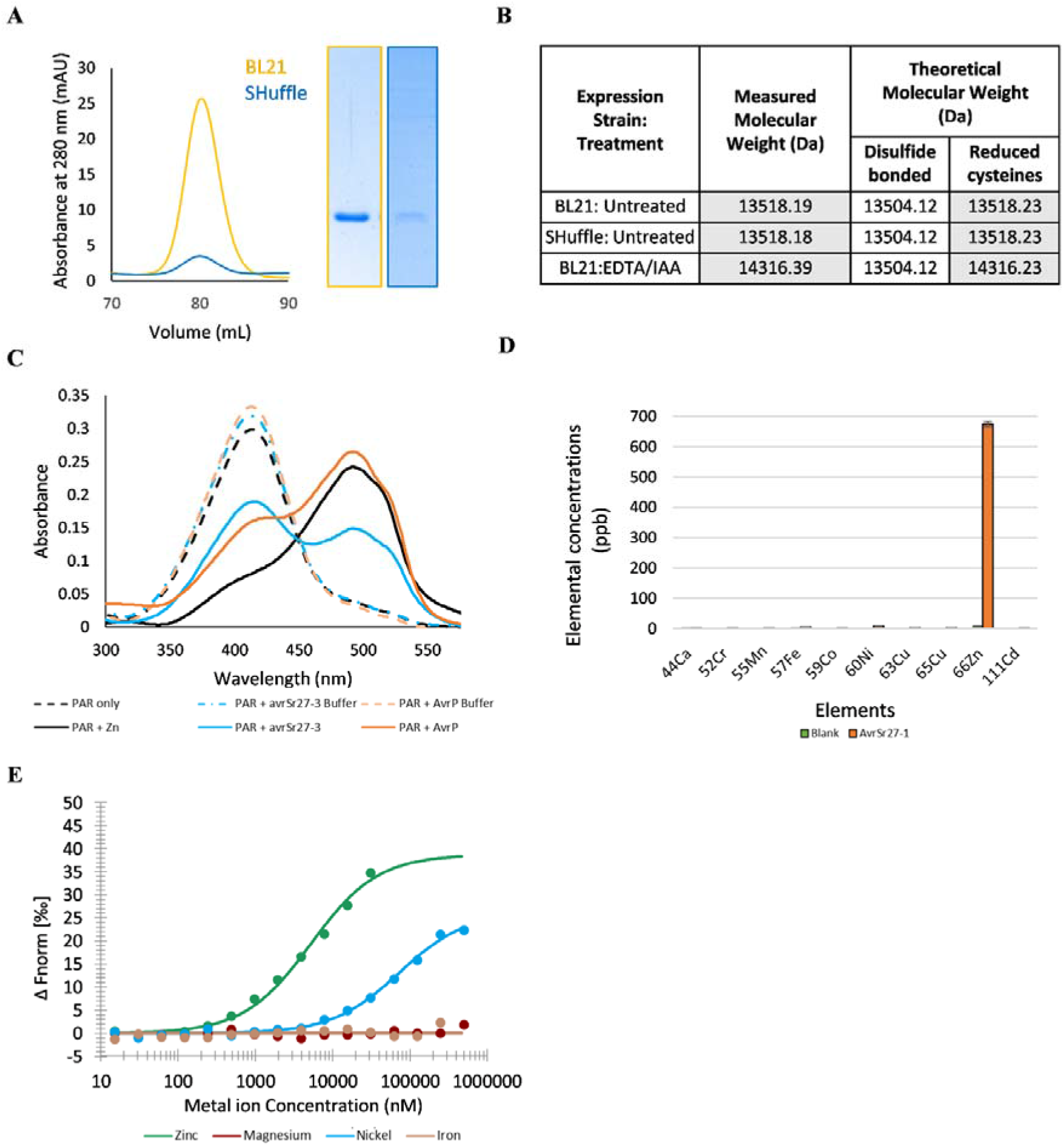
AvrSr27 is a metal-bound effector with a preference for zinc ions. (A) Size-exclusion chromatograms (SEC; left) using a HiLoad S75 16/600 column of nickel-affinity purified avrSr27-3 recombinantly produced in either the *E. coli* strain SHuffle or BL21. Coomassie-stained SDS-PAGE (right) of equal volume loading of fractions corresponding to the SEC peaks. **(B)** Intact mass analysis of avrSr27-3 protein produced either a reducing (BL21) or oxidising (SHuffle) environment. Proteins were either untreated or alkylated using iodoacetamine (IAA) after treatment with EDTA. The non-native residues that remained after 3C cleavage were included in this calculation. Shaded cells represent a match between the measured and theoretical molecular weight calculations. **(C)** Detection of metal ions bound in BL21 produced avrSr27-3 protein using 4LJ(2LJpyridylazo)resorcinol (PAR). The absorbance spectra are shown for PAR alone, PAR plus zinc ions, PAR plus the protein-less buffer controls for AvrP and avrSr27-3 and PAR plus denatured avrSr27-3 and AvrP, according to the key inset. **(D)** ICP-MS analysis of AvrSr27-1 (10 μM) diluted 1:1 with HNO3 (4 %v/v) overnight at room temperature. The results show the concentrations of calcium, chromium, manganese, iron, cobalt, nickel, copper, zinc, cadmium metal ions in the samples. Buffer indicates AvrSr27-1 SEC buffer. Error bars represent standard deviation across three independent experiments. **(E)** MST affinity measurements between fluorescently labelled AvrSr27-1 and transition metals ions. The affinity curves correspond to a Kd of 5.2 ± 0.74 μM for Zn^2+^(green) and 72.8 ± 7.2 μM for Ni ^2+^ (blue), while no binding was observed with Mg^2+^ (dark red) and Fe^2+^ (brown). Data points are calculated from three independent MST measurements.

To test if avrSr27-3 co-purified with metal ions, we expressed the protein in media supplemented with a trace metal solution and subsequently denatured the purified protein in the presence of the chromogenic chelator 4L(2Lpyridylazo) resorcinol (PAR). Free PAR has a maximum absorption wavelength of 416 nm, and shift to ∼500 nm (metal-dependent) when a PAR-metal ion complex is formed. After incubation of PAR with avrSr27-3 (denatured to release metal ions) a shift in the absorbance peak to ∼500 nm was observed, consistent with the formation of the PAR–metal complex (Fig. 1D). A similar shift was also observed upon incubation of PAR with AvrP, a known metal binding effector [19]. We confirmed the identity and stoichiometry of metal-binding in AvrSr27 using inductively coupled plasma mass spectroscopy (ICP-MS). AvrSr27 was found to be bound exclusively to zinc ions, with an average occupancy of 2.04 zinc ions per monomer of AvrSr27 (Fig 1D, Fig S3B). To evaluate the specificity and affinity of AvrSr27 for metals we used microscale thermophoresis (MST). We first sought to remove bound zinc from AvrSr27 by treating with EDTA under non-denaturing conditions, however subsequent analysis using the PAR assay and ICP-MS (post SEC) showed that AvrSr27 remained bound to zinc, with ratios similar to the untreated sample (∼1.7 zinc ions per AvrSr27 monomer) (Fig. S3A, B). Despite this, binding experiments with this protein using four transition metals (Zn^2+^, Mg^2+^, Ni^2+^, Fe^2+^) showed that the protein could still accommodate additional metals. The highest affinity was observed for Zn (Kd of 5.2 ± 0.74 μM) and a much lower affinity for Ni (Kd 72.8 ± 7.2 μM). No binding was observed for Mg^2+^ or Fe^2+^. Collectively, these data demonstrate that AvrSr27 can bind metals with a preference for zinc and can likely accommodate ≥2 Zn ions per molecule.

### The structure of AvrSr27 is a novel, duplicated domain

To elucidate the structure of AvrSr27, the three alleles were produced in *E. coli* BL21 and crystallisation screening was performed following a metal-exchange step to create a homogenous, zinc-loaded protein sample. Crystals of AvrSr27-1 were obtained, and the structure was solved to a final resolution of 2.4 Å using zinc single-wavelength anomalous diffraction (SAD) approach. The structure is comprised of two anti-parallel beta sheets, and a single short C-terminal helix (Fig. 2A). In the structure, there are four zinc ions, each bound with a tetrahedral coordination geometry by either four cysteine residues (CCCC; Zn2 and Zn4) or three cysteines and one histidine (CCCH; Zn1 and Zn3) (Fig 2B). The two beta sheet elements (residues 32-86, and 87-146, respectively), each with two Zn coordination sites, are structurally alike (root-mean squared deviation (RMSD) of 2.5 Å across 51 aligned residues; Fig. 2C), representing a duplicated domain with the second domain rotated 180 degrees with respect to the first. Each domain contains two zinc co-ordination sites with alternative co-ordination geometry (CCCH and CCCC). Interestingly, in the first zinc co-ordination site (Zn1), three cysteine residues (C37, C66 and C69) co-ordinate the zinc-ion together with histidine (H141), located in the C-terminal helix (Fig. 2B, C), which wraps around to stabilise the overall fold of the protein. Comparisons to the AlphaFold2 v2.3.0 predictions (Fig. S1B) showed that the predicted structures of AvrSr27-1 and avrSr27-3 showed resemblance to our experimentally determined structure (RMSD of ∼3 Å across 104, and 92 residues, respectively), but not AvrSr27-2 (Fig. S4A). The positions of the zinc co-ordinating residues in the AlphaFold2 (AF2) prediction of AvrSr27-1 were accurate except in the first binding site where the co-ordinating His residues differ (His33 vs His141 in the crystal structure) (Fig. S4B). Additionally, the avrSr27-3 AF2 structure predicted several disulfide bonds between cysteine residues in the Zn coordination sites (Fig. S4C).

**Figure 2:**
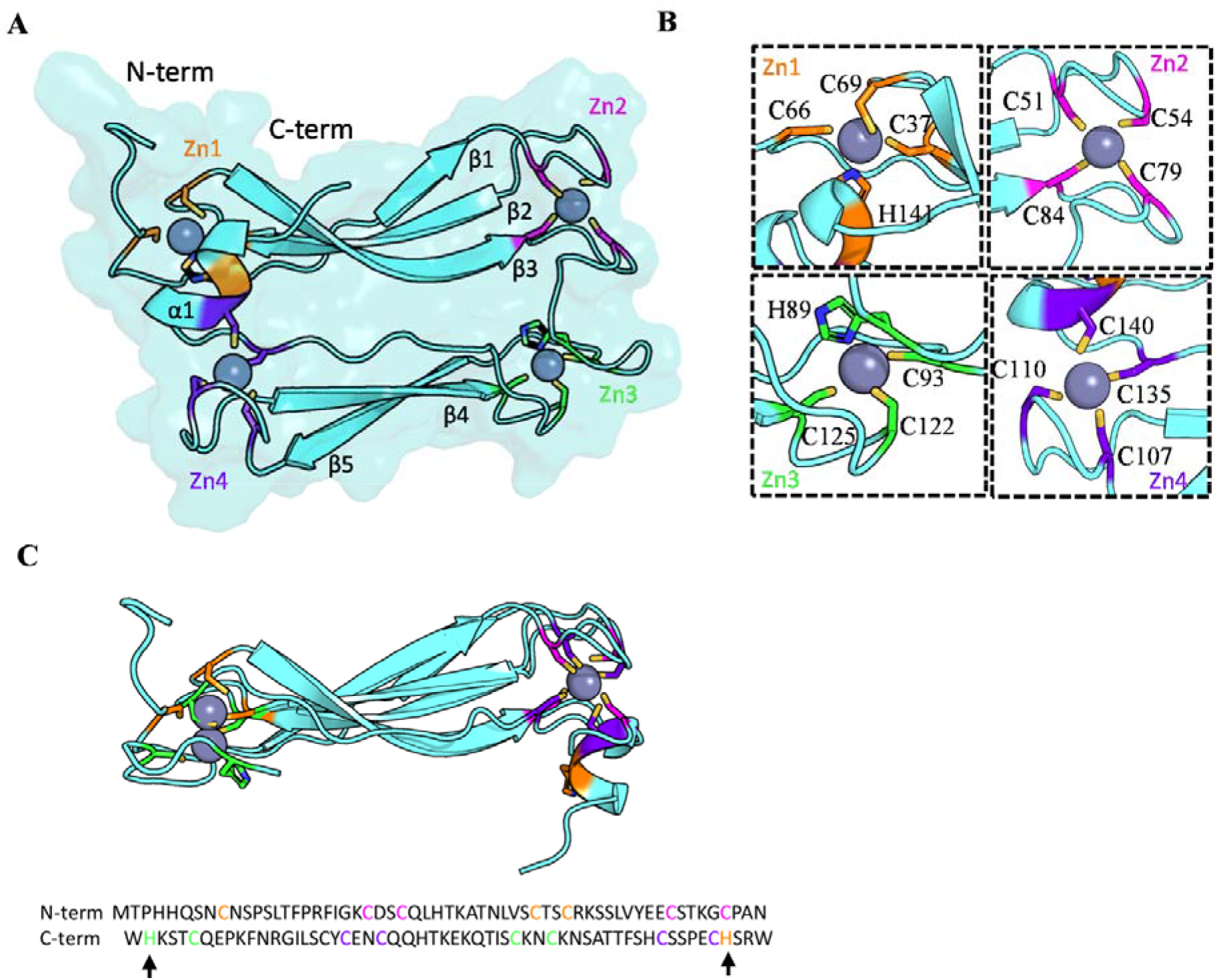
The crystal structure of AvrSr27-1 reveals a novel, metal-bound effector fold. (A) Cartoon representation of the overall structure of AvrSr27-1. Spheres represent the four bound Zn ions, with coordinating residues shown in stick form and coloured orange (Zn1 site), magenta (Zn2), green (Zn3) or purple (Zn4). **(B)** The tetrahedral coordination of Zn ions bound in the AvrSr27 structure with residue combinations (shown in stick form) of either CCCC for Zn2 and Zn4, and CCCH for Zn1 and Zn4. **(C)** Structural alignment of the AvrSr27-1 N-terminal domain and C-terminal domain (both in teal) shown in cartoon representation. Residues involved in Zn ion co-ordination are shown in stick form and coloured as above. Sequence alignment of the N- and C-term domains with co-ordinating Zn residues coloured as shown in the structure. The black arrows indicate the His residues involved in co-ordinating Zn1 and Zn3 pockets.

AvrSr27-1 has a novel structure amongst effectors, though limited structural similarity was observed to the substrate binding domain of the molecular chaperones, HSP70 and DnaK following a structural search of the protein databank (PDB) using the DALI web server [31] (Table S4, Fig. S5A,B). However, unlike the AvrSr27 proteins, these domains do not bind metal ions, nor does AvrSr27 contain the additional nucleotide-binding domain required for function in HSP70 and DnaK, suggesting it is unlikely to share the same functional role.

### Structure-informed identification of two proteins from *Pgt* that are recognised by and directly interact with Sr27

The two duplicated domains of AvrSr27 contain identical spacing between the Zn coordinating cysteine residues (Fig. 2C). A search for this motif (CX_13_CX_2_CX_11_CX_2_CX_9_CX_4_C; where X is any residue) identified two proteins from the *Pgt21-0* secretome, Pgt21-028479 and Pgt21-027343. These proteins share ∼95% sequence identity with one another (Fig. S6A), and ∼50% with the three AvrSr27 alleles (Fig. 3A) and are highly expressed in haustoria (Fig. S6B). Given proteins >30% sequence identity usually adopt a similar fold, we predict that Pgt21-028479 and Pgt21-027343 are structural homologues of AvrSr27.Intriguingly, transient co-expression of Pgt21-027343 with Sr27 in *N. benthamiana* resulted in a cell death response, that was slightly weaker than for AvrSr27-1, while Pgt21-028479 produced a quite weak response, indicating both proteins can be recognised by Sr27 (Fig. 3B, S7A, S8). Co-expression of AvrSr50 with Sr27 was included as a negative control (Fig. 3B-C, S7), while a lack of cell death observed for Pgt21-027343 or Pgt21-028479 when co-infiltrated with YFP, demonstrate the cell death is dependent on Sr27 recognition. Co-expression of Pgt21-027343 or Pgt21-028479 with Sr27 in wheat protoplasts also resulted in cell death as detected by a decrease in the percentage of living cells expressing the reporter YFP (Fig 3D). Expression of Pgt21-027343 or Pgt21-028479 alone showed no response, while there was also no response for co-expression of Sr27 with AvrSr50.

**Figure 3:**
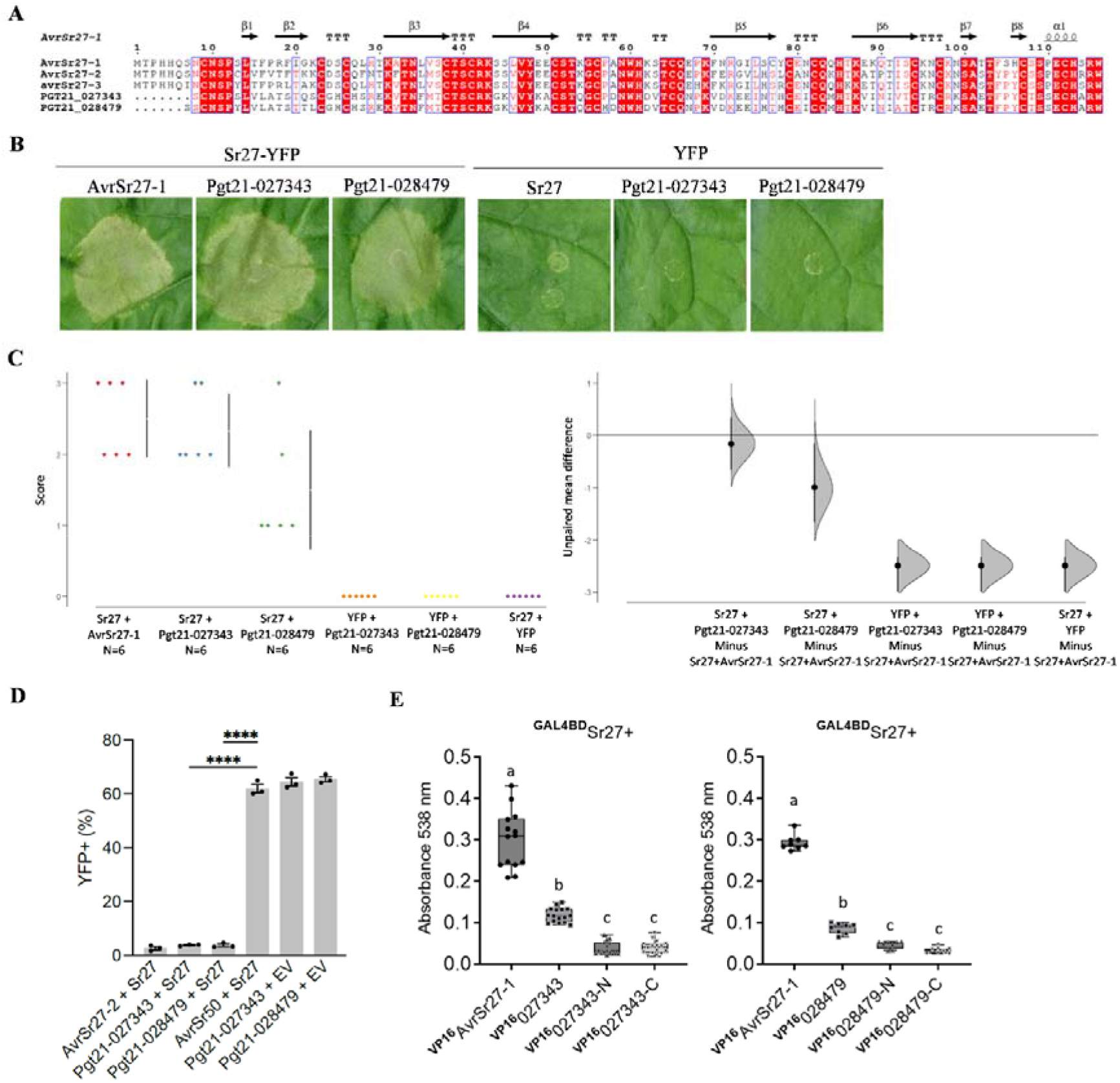
A Zn-binding motif searches uncovers two additional *Pgt* effectors that can be recognised by Sr27. (A) Multiple sequence alignment of the three AvrSr27 allelic and two motif identified proteins (Pgt21-028479 and Pgt-027434) generated using ESPript (Robert & Gouet, 2014 [38]). The secondary structure elements of our experimentally determined crystal structure (AvrSr27-1) are displayed on the alignment. ‘T’ corresponds to a β-turn. **(B)** Representative cell death response in *N. benthamiana* leaves when transiently co-expressing YFP-tagged AvrSr27-1, Pgt21-028479, Pgt21-027343 with pAM-GWY-Sr27-YFP or YFP alone **(C)** Cummings estimation plot showing cell death phenotypes as represented in B and Fig. S7. The left panel show the distribution of the observed scores for each protein where each dot represents an independent infiltration assay and the number of replicates per sample are indicated on the x-axis. The right panel represents the mean differences for comparisons of variants against a shared control (AvrSr27-1 + Sr27). **(D)** Flow cytometry assay for cell death in transiently transformed wheat protoplasts. The percentage of living cells expressing a YFP reporter is shown after co-transformation with combinations of Sr27 or an empty vector (EV) with AvrSr27-2, Pgt21-027343, Pgt21-028479, or AvrSr50 Error bars represent the standard error of the mean (n = 3). ****, p < 0.0001 based on a upaired t-test assuming equal variances. **(E)** Quantification of betalain accumulation by absorbance at 538 nm for leaf infiltration sites expressing GAL4BD-Sr27 with VP16-fused to Pgt21-027343 (left graph), Pgt21-028749 (right graph) and their N and C terminal domains respectively. Common letters above columns indicate no significant difference between samples (p>0.05; one way ANOVA with posthoc Tukey HSD). Further replicates are shown in Fig. S9 and Fig. S8.

We recently showed that Sr27 physically associates with AvrSr27-1, -2 and-3 using a plant two-hybrid assay in which production of the purple pigment betalain by a *Ruby* reporter gene serves as a readout for protein-protein interaction [16]. Co-expression of GAL4BD-Sr27 with VP16-fused Pgt21-028479 and Pgt21-027343 resulted in activation of the UAS (upstream activating sequence)-Ruby reporter and production of betalain (Fig 3E, S8, S9), indicating physical association between these effector proteins and Sr27. However, betalain accumulation was significantly less than that observed for the Sr27-AvrSr27-1 positive control (Fig. 3E, S8-9) despite the proteins accumulating to similar levels as detected by immunoblotting (Fig S10), suggesting that the Pgt21-028479/Sr27 and Pgt21-27343/Sr27 associations are weaker.

### The N-terminal domain of AvrSr27 is sufficient for recognition and association with Sr27

To investigate the requirements for the interaction between Sr27 and the AvrSr27 and AvrSr27-like proteins we split the proteins into their N- and C-terminal domains (Table S2). For the three AvrSr27 alleles an additional shorter construct (residues 35-86, termed AvrSr27-N-short), was created that excludes the two N-terminal His residues, which could potentially complete the first Zn co-ordination site in the absence of His141 in these fragments (Fig. 2). Following transient co-expression with Sr27 in *N. benthamiana* and *N. tabacum*, a strong cell death response was observed for all the AvrSr27 N-terminal constructs, similar to the full-length AvrSr27 proteins (Fig. 4A, S11-14). However, little or no cell death was observed for the AvrSr27-1 and AvrSr27-2 C-terminal domains, while only a weak response was observed for avrSr27-3 (Fig. 4A, Fig. S13). Unlike the AvrSr27 alleles, no cell death was observed for either the N- or C-terminal constructs of Pgt21-027343 and Pgt21-028479, nor could recognition of these protein fragments be recovered when both N- and C- constructs were co-expressed together with Sr27 (Fig. 4B, S14).

**Figure 4:**
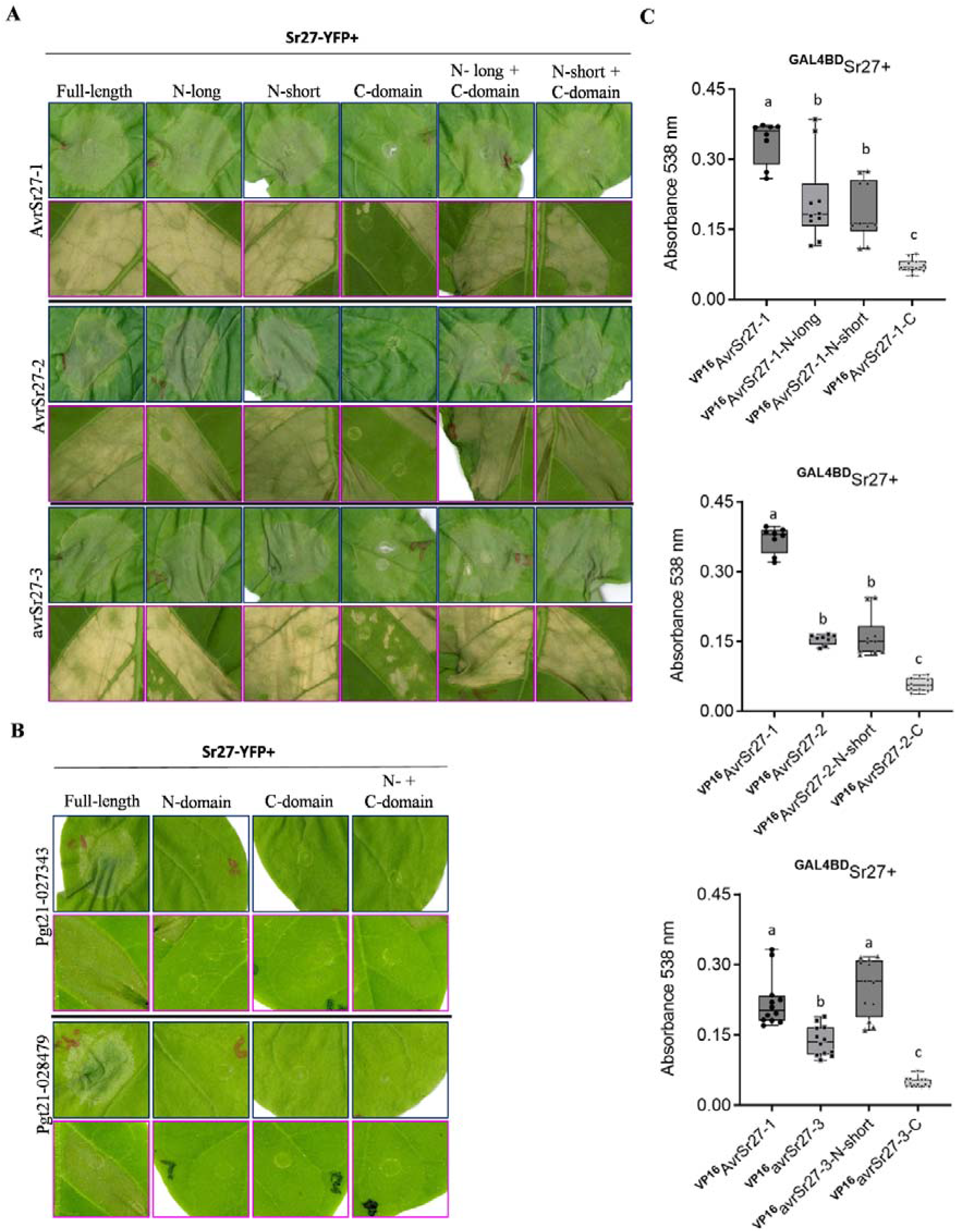
The N-terminus of AvrSr27 is sufficient to confer recognition and direct interaction with Sr27. Representative *N. benthamiana* (dark blue outline) and *N. tabaccum* (cyan outline) leaf infiltration sites showing cell death following transient co-expression of Sr27 with **(A)** full length and N- or C-terminal fragments of AvrSr27 constructs and **(B)** full length and N- or C-terminal fragments of Pgt21-027343 or Pgt21-028479. All leaves and cumming estimation plots (for AvrSr27 proteins only) are shown in Fig S11-14. **(C)** Quantification of betalain accumulation by absorbance at 538 nm for leaf infiltration sites expressing GAL4BD-Sr27 with VP16 fused to AvrSr27-1, -2, -3 and their N- and C terminal domains. Common letters above columns indicate no significant difference between samples (p>0.05; one way ANOVA with posthoc Tukey HSD). Additional experiments and corresponding leaves are shown in Fig S15 and S16.

To investigate if the Sr27 physical association was maintained for the AvrSr27 N-terminal domain we carried out the *in planta* two-hybrid assay as before and assessed betalain production. For AvrSr27-1, the N-terminal long and short fragments gave betalain accumulation when co-expressed with GAL4BD-Sr27, but slightly less than that for the full length protein (Fig. 4C, S15A, S16A), while the N-terminal (short) fragments of AvrSr27-2 gave similar levels to the full length proteins (Fig. 4C, S15B, S16B) and the avrSr27-3 N-terminal (short) domain showed a similar interaction to the full length protein. Little to no betalain accumulated following co-infiltration of the AvrSr27 C-terminal domains, nor the Pgt21-028479 and Pgt21-027343 N- or C-domain splits (Fig 3C, S8-S9). Protein expression levels for all constructs was similar (Fig S8A).

## Discussion

Determining the molecular basis of effector recognition by host immune receptors is important for understanding the evolution of virulence traits in pathogen populations and can underpin the successful deployment of existing genetic resistance against phytopathogens, such as *Pgt* [39]. Recognition between the wheat immune receptors Sr27, Sr35 and Sr50 and their corresponding Avr proteins from *Pgt* is mediated by direct protein interactions [6, 16, 40]. We previously found that a mutation in a single amino acid surface exposed residue was sufficient for a virulence allele of AvrSr50 to escape recognition by the Sr50 immune receptor, and defined part of the interaction surface [12]. The Sr35-AvrSr35 resistosome structure identified an interaction site between these proteins involving mainly the 10^th^ alpha helix of AvrSr35 contacting the latter half of the LRR domain of Sr35, with residues within this region confirmed to be important for binding and activation in functional assays [14, 15]. In this study, we demonstrate biochemically and structurally that AvrSr27 is a zinc-binding effector with a novel modular fold consisting of two structurally similar domains. The N-terminal domain of AvrSr27 is sufficient for recognition and interaction with the corresponding wheat Sr27 immune receptor. Further, using a structurally identified metal-binding motif we identified two additional putative *Pgt* effectors that can also be recognised by Sr27, though only as full-length proteins.

### AvrSr27 is a zinc-bound effector

Many candidate effector proteins from filamentous phytopathogens are enriched for cysteine residues, which are often considered to be involved in intra- and intermolecular disulfide bonds, providing structural stability [19]. Largely, this has held true for effector proteins that localise to the apoplast. Conversely, effectors delivered to the cytosol tend to be low in disulfides (and cysteine content), and while there are examples of disulfide bond containing cytoplasmic effectors (e.g. MAX effectors from *Magnaporthe oryzae* and ToxA from *Pyrenophora tritici-repentis*), these effectors are not necessary cysteine rich. Prior to this study, the structural characterisation of two cytoplasmic-localising cys-rich effectors (AvrP from the flax rust fungus *M. lini* [17], and AvrPii from the rice blast fungus *Magnaporthe oryzae* [18]) demonstrated that the cysteines were involved in coordinating zinc-ions. Here, we showed that mature AvrSr27, which is ∼12% cysteine, binds to 4 zinc ions, coordinated by its cysteine residues. The highly sensitive ICP-MS approach demonstrated that the protein preferentially binds zinc ions from a mixed solution of metal ions. This approach provides a useful addition to our previously described workflow [19] to understand whether cys-rich effector proteins are adopting disulfide-bonded or metal bound structural states. We attempted to understand the affinity of AvrSr27 for zinc (following EDTA treatment) using MST however, both the PAR and ICP-MS experiments showed that AvrSr27 could not be completely stripped of Zn. Based on our structure, ICP-MS and MST data, we speculate that the EDTA treatment removes two of the four zinc ions bound to the protein. Collectively, these data suggest AvrSr27 contains two high and two weaker affinity binding sites (those that we could measure via MST), however the biological significance of this is yet to be realised.

The observation that AvrSr27, AvrP and AvrPii all bind metal ions via cysteine coordination, suggests that this may be a common feature of cys-rich cytoplasmic effectors, but the function(s) of this remain unclear. Binding of zinc and other metal ions are known to play a structural role ensuring protein stability and the induction of correct folding, which in lieu of disulfide bonds may provide rigidity and protection following secretion into the plant cell. However, zinc has also been shown to be essential for catalytic activities in many enzymes. Experimental evidence suggests that the functionality of Zn ions within a protein is dependent on the coordination environment [41, 42] and the tetrahedral (CCCC and CCCH) coordination of Zn observed in the structures of these effectors is more representative of a structural role. Another intriguing possibility is that during infection metal binding effectors may be produced in the fungus without bound Zn and may then act to sequester Zn within the plant cell after delivery to the host. Zinc ions, and other transition metals play important roles in plant growth and physiology, including in responses to either abiotic or biotic stresses, and their intracellular concentrations are tightly regulated to maintain biological processes [43]. High local concentrations of Zn are reportedly toxic to bacteria and fungi, and in response to pathogen attack plants can raise the local zinc concentration to either create a toxic, inhospitable environment, or activate other plant defence processes [43]. For example, zinc is reported to induce oxidative stress and the generation of reactive oxygen species (ROS), which is effective against biotrophic pathogens like *Pgt* [44].

### AlphaFold2 capacity to predict effector proteins shows varied results

In the last couple of years there has been significant advancements provided by AI-based structural predictors, such as AlphaFold2 [34], and has proven to be extremely useful in effector studies [45-52]. Despite this, accuracy for effector prediction (compared to experimental structures) remain varied [47, 53, 54]. A factor that may contribute to this is the evolutionary depth of the multiple sequence alignments [47, 53], which underpin AlphaFold2, although in some cases it remains unclear. For AvrSr27 (variants and structural homologues) the predictions were generally poor, and varied despite high sequence conservation (Fig. S1A, S6C). We suggest that the contribution of zinc as a co-factor for AvrSr27 and the current inability for deep learning structural prediction algorithms to incorporate co-factors during the prediction process may be the reason for this. Though we note that AlphaFold2 has shown capacity to correctly predict zinc co-ordination sites in some instances [34]

### Modularity in effector proteins

AvrSr27 consists of two structurally related domains repeated in tandem, which is reminiscent of the conserved structural modules described in effectors of some other phytopathogens, although not previously reported in rust pathogens. For example, the transcription activator-like effectors (TAL) produced by *Xanthomonas* bacterial species contain modular tandem repeats with variant residues that determine DNA binding specificity of these effectors [55], which act as transcription factors that change expression of targeted genes to promote infection [56]. Likewise, hundreds of RXLR effectors from *Phytophthora spp.* contain various numbers of the three or four α-helix bundle WY domain. For instance, the WY effector *Ps*PSR2 (*P. sojae* Phytophthora suppressor of RNA silencing 2) contains seven tandem repeats where the first module is a canonical WY domain, and the other six repeats contain a LWY motif and a highly conserved fold consisting of five α-helices [57]. The different molecular functions established for various WY effectors suggest that their modular structure supports the diversification of new virulence functions [58-61]. Recently, a functional L(WY)-LWY module was identified in 12 effectors from two *Phytophthora* species that recruit and hijack a Serine/Threonine protein phosphatase 2A (PP2A) core enzyme, and through diversification in the C-terminal LWY module alters the specificity of targeted host phosphotases [62]. Here, we observed that AvrSr27 consists of two repeated domains, indicating that *Pgt* effectors, like *Phytophthora* and bacterial pathogens, likely utilise core structural modules.

### Structural insights into effector recognition by Sr27

The modular structure of AvrSr27 has implications for the recognition by the host Sr27 immune receptor. The N-terminal domain of AvrSr27 is sufficient for recognition and direct interaction with Sr27, while the C-terminal domain is not recognised. Unexpectedly two AvrSr27-like structural homologues (Pgt21-027343 and Pgt21-028479) could also interact with Sr27 and induce cell death when co-expressed as full-length proteins, but neither their N- nor C-terminal domains are recognised. This clear difference suggests that there are conserved surface exposed residues involved in mediating this interaction in the AvrSr27 alleles that differ in the AvrSr27-like structural homologues. Pgt21-027343 and Pgt21-028479 show high expression in haustoria and during infection but do not confer an avirulence phenotype, since mutants virulent on Sr27 retain these genes intact [7]. This failure to trigger resistance during infection may be due to the weaker association between these proteins and Sr27 as compared to the AvrSr27 variants (Fig. 3B, C). Alternatively, although transcribed in the fungus, these genes may not be efficiently translated, or the translated products may not be delivered to the host. Another possibility is that these dual-domain proteins are processed during infection to single domains, which is consistent with the observation that Sr27 can recognise the N-terminal domains of the AvrSr27 proteins but not those of Pgt21-027343 and Pgt21-028479 (Fig. 4A, B). Along with the previously observed *in planta* recognition of the protein encoded by the low-expressed *avrSr27-3* virulence allele [7], these data suggest that recognition between co-expressed R and Avr proteins may not always correlate with resistance/virulence phenotypes during infection, and as such caution is warranted in interpreting such experiments in the absence of infection phenotype data.

## Acknowledgements

The work described here was supported in part by funding from the 2Blades Foundation. SW was supported by an Australian Research Council Future Fellowship (FT200100135). MO was supported by an AINSE Early Career Research Grant. MO and JC are supported by CSIRO Research Postdoctoral Fellowships. The authors acknowledge the use of the ANU Crystallisation and X-ray facility. The authors also acknowledge use of the Australian Synchrotron MX facility and thank the staff for their support. Additionally, the team acknowldges use of the CSIRO High Performance Computing Facility and expertise provided by the CSIRO IMT Scientific Computing and Data61 teams.

## Author contributions

MO, JC, PD, and SW designed the study; MO, JC, SB, TA, CB, ZL, DE, JS, MF, NT, and SA performed the experiments and all authors analysed the data. MO, JC, SW, and PD wrote the original manuscript draft, and all authors contributed to writing, reviewing, and editing the manuscript.

## Data availability

The structure of AvrSr27 has been deposited to the protein databank under the PDB code 8V1J.

