## supplementary figures and tables for "AvrSr27 is a zinc-bound effector with a modular structure important for immune recognition"

The following Supporting Information is available for this article:

Tables S1 to S3.

Figures S1 to S16.

SI References.

**Table S1 Primers used in this study.**

| **Primer name** | **Primer sequence (5'-3')** |
| --- | --- |
| AvrSr27-1 and AvrSr27-2 GG Fw | TAGGTCTCCAATGACACCACATCACCAAAGCA |
| AvrSr27-3 GG Fw | TAGGTCTCCAATGACACCACATCACCAAATCAATT |
| AvrSr27-1 and Avrsr27-3 GG Rv | ACGGTCTCCAAGACCATCTGCTGTGACACTCTGG |
| AvrSr27-2 GG Rv | ACGGTCTCCAAGACCATCTTCTGTGACACTCTGGG |
| PGT21_027343 GG Fw | TAGGTCTCCAATGAGCAGCTGCAACTCACCATC |
| PGT21_028479 GG Fw | TAGGTCTCCAATGAGCAGCTGCAACTCACCATAT |
| PGT21_027343_028479 GG Rv | ACGGTCTCCAAGACCATCTTGCATGACACTCTGAGC |
| AvrSr27_C1 N-term GG Rv | ACGGTCTCCAAGAAGCAGGACAGCCTTTGGTG |
| AvrSr27_Cterm GG Fw | TAGGTCTCCAATGAATTGGCACAAAAGCACCTGTCA |
| AvrSr27_Cterm GG Rv | ACGGTCTCCAAGACCATCTGCTGTGACACTCTGGG |
| AttB1-AvrSr27C1/2-d28 | GGGGACAAGTTTGTACAAAAAAGCAGGCTTAATGACACCACATCACCAAAGCAAT |
| AttB1-AvrSr27C3-d28 | GGGGACAAGTTTGTACAAAAAAGCAGGCTTAATGACACCACATCACCAAATCAAT |
| AttB1-AvrSr27C1-d34 | GGGGACAAGTTTGTACAAAAAAGCAGGCTTAATGAGCAATTGCAACTCCCCATCTTTGA |
| AttB1-AvrSr27C2-d34 | GGGGACAAGTTTGTACAAAAAAGCAGGCTTAATGAGCAATTGCAACTCCCCATGTTTGG |
| AttB1-AvrSr27C3-d34 | GGGGACAAGTTTGTACAAAAAAGCAGGCTTAATGATCAATTGCAACTCCCCATATTTGACA |
| AttB2-AvrSr27C1/2/3-66R | GGGGACCACTTTGTACAAGAAAGCTGGGTCTTAAGCAGGACAGCCTTTGGTGG |
| AttB1-AvrSr27C1-67 | GGGGACAAGTTTGTACAAAAAAGCAGGCTTAATGAATTGGCACAAAAGCACCTGTCAA |
| AttB1-AvrSr27C2-67 | GGGGACAAGTTTGTACAAAAAAGCAGGCTTAATGAATTGGCACAAAAGTACCTGTCAA |
| AttB1-AvrSr27C3-67 | GGGGACAAGTTTGTACAAAAAAGCAGGCTTAATGAATTGGCACAAAAGTACCTGTCAA |
| AttB2-AvrSr27C1/3 | GGGGACCACTTTGTACAAGAAAGCTGGGTCTTACCATCTGCTGTGACACTCTGG |
| AttB2-AvrSr27C2 | GGGGACCACTTTGTACAAGAAAGCTGGGTCTTACCATCTTCTGTGACACTCTGG |
| AttB1-AvrSr27-027343/028479 | GGGGACAAGTTTGTACAAAAAAGCAGGCTTAATGAGCAGCTGCAACTCACCA |
| AttB2-AvrSr27-027343-53R | GGGGACCACTTTGTACAAGAAAGCTGGGTCATCAGGACAGCCTTGGGTG |
| AttB2-AvrSr27-028479-53R | GGGGACCACTTTGTACAAGAAAGCTGGGTCATCATGACAGCCTTGGGTGGA |
| AttB1-AvrSr27-027343/028479-54F | GGGGACAAGTTTGTACAAAAAAGCAGGCTTAATGAATTGGCATGATGTCACTTGTCA |
| AttB2-AvrSr27-027343/028479 | GGGGACCACTTTGTACAAGAAAGCTGGGTCCCATCTTGCATGACACTCTGAG |

**Table S2: List of constructs used in this study.**

| **Construct name** | **Insert or PCR product** | **Backbone** | **Primers** |
| --- | --- | --- | --- |
| 6xHis-3C-AvrSr27-1 | AvrSr27-1 29-144 | pOPIN_F3_RFP_AmpR | AvrSr27-1 GG Fw and AvrSr27-GG Rv |
| 6xHis-3C-AvrSr27-2 | AvrSr27-2 29-144 | pOPIN_F3_RFP_AmpR | AvrSr27-2 GG Fw & AvrSr27-GG Rv |
| 6xHis-3C-AvrSr27-3 | AvrSr27-3 29-144 | pOPIN_F3_RFP_AmpR | AvrSr27-3 GG Fw & avrsr27-1 GG Rv |
| Sr27-YFP | Sr27 | pAM-GWY-YFP | (Upadhyaya et al., 2021) |
| YFP-AvrSr27-1 | AvrSr27-1 29-144 | pBIN19-YFP-GWY |  |
| YFP-AvrSr27-2 | AvrSr27-2 29-144 | pBIN19-YFP-GWY |  |
| YFP-AvrSr27-3 | AvrSr27-3 29-144 | pBIN19-YFP-GWY |  |
| YFP-N-long-AvrSr27-1 | AvrSr27-1 29-87 | pBIN19-YFP-GWY | AttB1-AvrSr27-1/2-d28 and AttB2-AvrSr27-1/2/3-66R |
| YFP-N-short-AvrSr27-1 | AvrSr27-1 34-87 | pBIN19-YFP-GWY | AttB1-AvrSr27-1-d34 and AttB2-AvrSr27-1/2/3-66R |
| YFP-N-long-AvrSr27-2 | AvrSr27-2 29-87 | pBIN19-YFP-GWY | AttB1-AvrSr27-1/2-d28 and AttB2-AvrSr27-1/2/3-66R |
| YFP-N-short-AvrSr27-2 | AvrSr27-2 34-87 | pBIN19-YFP-GWY | AttB1-AvrSr27-2-d34 and AttB2-AvrSr27-1/2/3-66R |
| YFP-N-long-AvrSr27-3 | AvrSr27-3 29-87 | pBIN19-YFP-GWY | AttB1-AvrSr27-3-d28 and AttB2-AvrSr27-1/2/3-66R |
| YFP-N-short-AvrSr27-3 | AvrSr27-3 34-87 | pBIN19-YFP-GWY | AttB1-AvrSr27-3-d34 and AttB2-AvrSr27-1/2/3-66R |
| YFP-Cterminal-AvrSr27-1 | AvrSr27-1 88-144 | pBIN19-YFP-GWY | AttB1-AvrSr27-1-67 and AttB2-AvrSr27-1/3 |
| YFP-Cterminal-AvrSr27-2 | AvrSr27-2 88-144 | pBIN19-YFP-GWY | AttB1-AvrSr27-2-67 and AttB2-AvrSr27-2 |
| YFP-Cterminal-AvrSr27-3 | AvrSr27-3 88-144 | pBIN19-YFP-GWY | AttB1-AvrSr27-3-67 and AttB2-AvrSr27-1/3 |
| YFP-027343 | Pgt21-027343 22-130 | pBIN19-YFP-GWY | Synthesised pDONR construct |
| YFP-028479 | Pgt21-028479 22-130 | pBIN19-YFP-GWY | Synthesised pDONR construct |
| YFP-Nterminal-027343 | Pgt21-027343 22-73 | pBIN19-YFP-GWY | AttB1-AvrSr27-027343/028479 and AttB2-AvrSr27-027343-53R |
| YFP-Nterminal-028479 | Pgt21-028479 22-73 | pBIN19-YFP-GWY | AttB1-AvrSr27-027343/028479 and AttB2-AvrSr27-028479-53R |
| YFP-Cterminal-027343 | Pgt21-027343 74-130 | pBIN19-YFP-GWY | AttB1-AvrSr27-027343/028479-54F and AttB2-AvrSr27-027343/028479 |
| YFP-Cterminal-028479 | Pgt21-028479 74-130 | pBIN19-YFP-GWY | AttB1-AvrSr27-027343/028479-54F and AttB2-AvrSr27-027343/028479 |
| pTA22 | NA | pTA22 | (Taj et al., 2023) |
| pTA22-YFP | Venus YFP | pTA22 |  |
| pTA22-Sr27 | Sr27 | pTA22-GW |  |
| pTA22-AvrSr27-2-PBS | AvrSr27-2 | pTA22-GW-PBS |  |
| pTA22-AvrSr50-PBS | AvrSr50 | pTA22-GW-PBS |  |
| pTA22-0285-PBS | Pgt21-027343 | pTA22-GW-PBS |  |
| pTA22-0286-PBS | Pgt21-028479 | pTA22-GW-PBS |  |

**Table S3: X-ray data collection, structure solution and refinement statistics for AvrSr27-1.**

|  | High remote (SAD) |
| --- | --- |
| **Data collection** |  |
| Detector | Eiger |
| Wavelength () | 1.26 |
| Crystal-to-detector distance (mm) | 200 |
| Space group | P 1 21 1 |
| *a*, *b*, *c* (Å) | 40.65 34.30 70.74 |
| a, b, g (°) | 90.00 92.90 90.00 |
| Average mosaicity (°)^b^ | 0 |
| Resolution (Å) | 43.59-2.42 (2.51-2.42) |
| Total no. of reflections | 128832 (12519) |
| No. of unique reflections | 8197 (811) |
| Completeness (%) | 99.2 (95.5) |
| Multiplicity | 15.7 (15.4) |
| Anomalous completeness | 98.6 (92.2) |
| Anomalous multiplicity | 8.0 (8.0) |
| Mean *I* /s(*I*) | 11.0 (3.9) |
| *R*meas (%)^c^ | 16.9 (58.3) |
| *R*pim (%)^d^ | 5.8 (20.0) |
| CC_1/2_^b^ | 0.997 (0.935) |
| Matthews coefficient (Å^3^ Da^-1^)^e^ | 1.96 |
| **Phasing statistics determined by Crank2** |  |
| No. of sites identified | 8 |
| Overall figure of merit (FOM) | 0.3525 |
| FOM after density modification | 0.5026 |
| Resolution range (Å) | 43.59-2.42 |
| *R*_work_ (%)^g^ | 19.3 |
| *R*_free_ (%)^h^ | 25.1 |
| No. of non-H atoms |  |
| Total | 3452 |
| Non-solvent | 3432 |
| Ligand | 8 |
| Water | 12 |
| Average *B*-factor (Å^2^) | 34.0 |
| R.m.s.d. from ideal geometry |  |
| Bond lengths (Å) | 0.006 |
| Bond angles (°) | 0.520 |
| Ramachandran plot, residues in (%)^i^ |  |
| Favoured regions | 96.0 |
| Additionally allowed regions | 4.0 |
| Outlier regions | 0.0 |

^a^ The values in parentheses are for the highest-resolution shell.

^b^ Calculated with AIMLESS.

^c^ *R*meas = Σ*_hkl_*{*N*(*hkl*)/[*N*(*hkl*)-1]}^1/2^ Σ*i*|I*i*(*hkl*)-<*I*(*hkl*)>|/ Σ*hkl*Σ*i*I*i*(*hkl*), where *Ii*(*hkl*) is the intensity of the *i*th measurement of an equivalent reflection with indices *hkl*.

^d^ *R*pim = Σ*_hkl_*{1/[*N*(*hkl*)-1]}^1/2^ Σ*i*|I*i*(*hkl*)-<*I*(*hkl*)>|/ Σ*hkl*Σ*i*I*i*(*hkl*).

^e^ Calculated with MATTHEWS_COEF within the CCP4 suite.

^f^ Generated by Crank pipeline in the CCP4 suite.

^g^ *R*_work_  = Σ*_hkl_*  ||*F*_obs_ |-|*F*_calc_||/Σ_hkl_ |*F*_obs_|, where *F*_obs_ and *F*_calc_ are the observed and calculated structure factor amplitudes.

^h^ *R*_free_  is equivalent to *R*_work_  but calculated with reflections (10%) omitted from the refinement process.

^i^ Calculated with MolProbity.

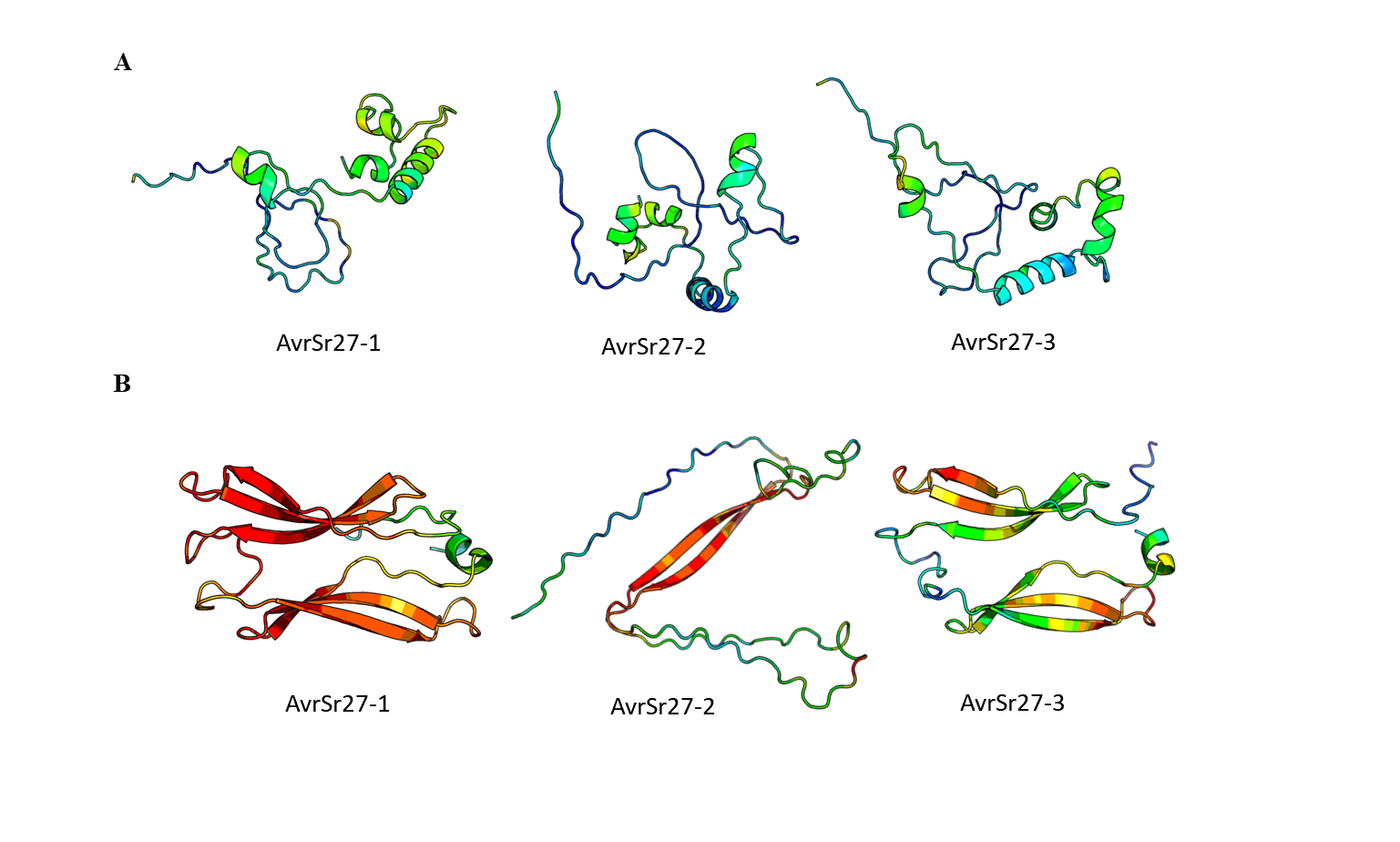

**Fig S1 AlphaFold2 predictions of the three AvrSr27 variants.** AlphaFold2 predictions obtained from (A) the AlphaFold2 database (<https://alphafold.ebi.ac.uk/>) and (B) using a local install of AlphaFold2 (as described in the methods and materials) for the three allelic variants of AvrSr27, from left to right, AvrSr27-1, -2, and avrSr27-3, represented in cartoon form. Models are coloured by predicted confidence scores (pLDDT), where red is most confident, and blue is least confident.

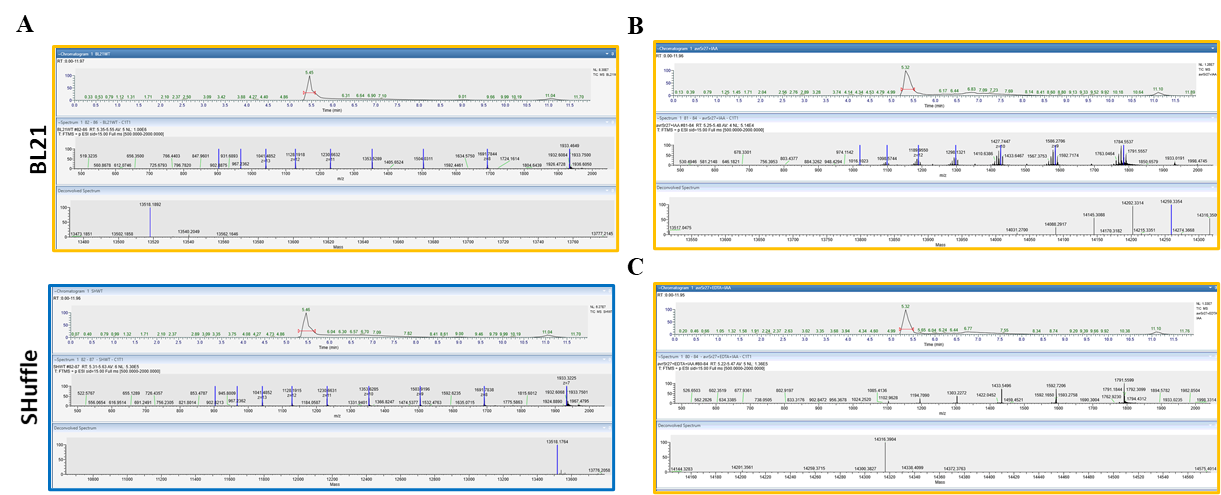

**Figure S2. Intact Mass Spectrometry spectra** for avrSr27-C3 proteins produced recombinantly in *E. coli* SHuffle (oxidising, blue) or BL21 (reducing, yellow) as shown in Fig. 2A and 2B. Proteins treatments are as follows **(A)** Untreated **(B)** Alkylated **(C)** EDTA/Alkylated avrSr27-3.

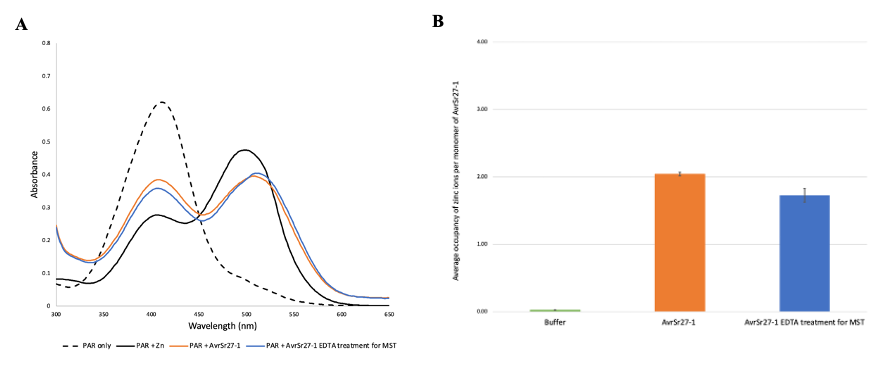

**Fig S3. AvrSr27-1 binds Zinc ions in PAR assay and ICP-MS.**

**(A)** Detection of metal ions bound in AvrSr27-1 protein using 4‐(2‐pyridylazo) resorcinol (PAR). The absorbance spectra are shown for PAR alone, PAR plus zinc ions, PAR plus AvrSr27-1 protein and PAR plus AvrSr27-1 protein treated with EDTA, according to the key inset. **(B)** The average occupancy of zinc ions per AvrSr27-1 monomer was calculated using ICP-MS. The orange column indicates AvrSr27-1 samples without treatment, the blue column indicates AvrSr27-1 samples treated with EDTA. Buffer indicates the AvrSr27-1 SEC buffer.

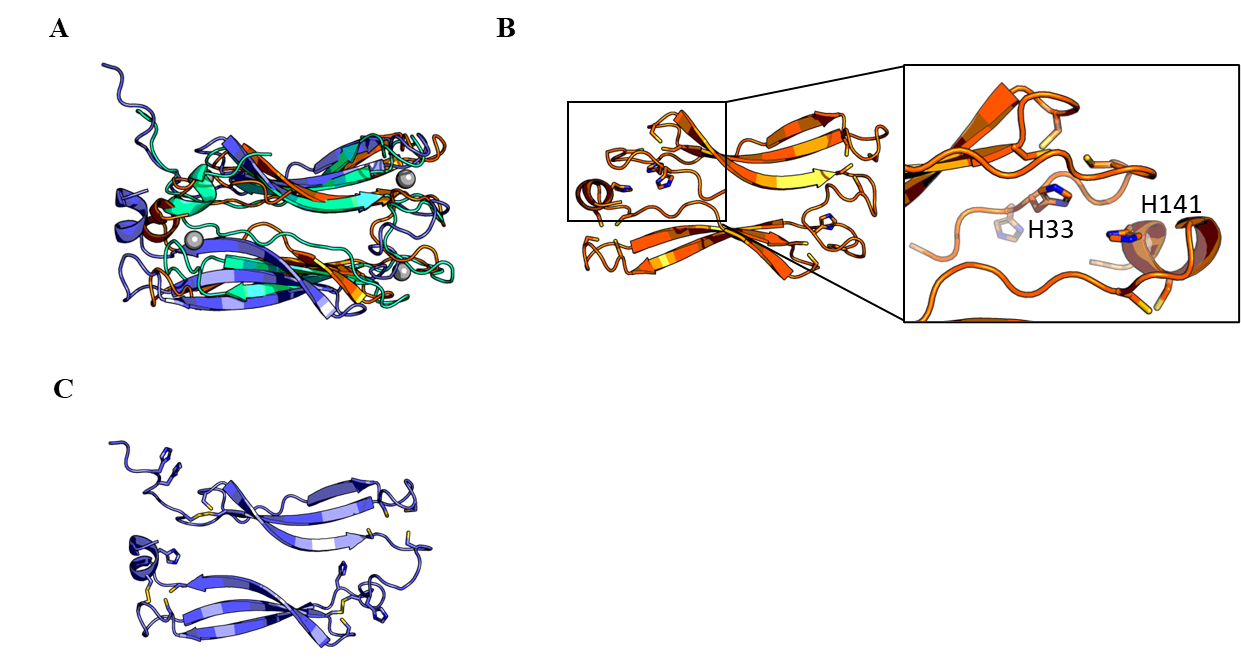

**Fig S4 Benchmarking AlphaFold2 against our experimentally determined structure of AvrSr27.** (A) Dali pairwise aligned structure superimposition of AvrSr27-1 experimental structure (teal), with the AlphaFold predictions of AvrSr21-1 (orange) and avrSr27-3 (purple) (shown in Fig S1B) from a local Alphafold2 (v2.3.0 install). Structures are all shown as cartoon representation. (B) AvrSr27-1 AlphaFold2 predicted structure showing the Zn co-ordinating residues in stick representation, and insert highlighting difference in His residue co-ordinating between our experimental structure (H141) and what is observed in the predicted structure (H33). (C) The avrSr27-3 AlphaFold2 predicted structure showing the Zn co-ordinating residues in stick representation. In three of the pockets two of the co-ordinating Cys residues are predicted to form disulfides.

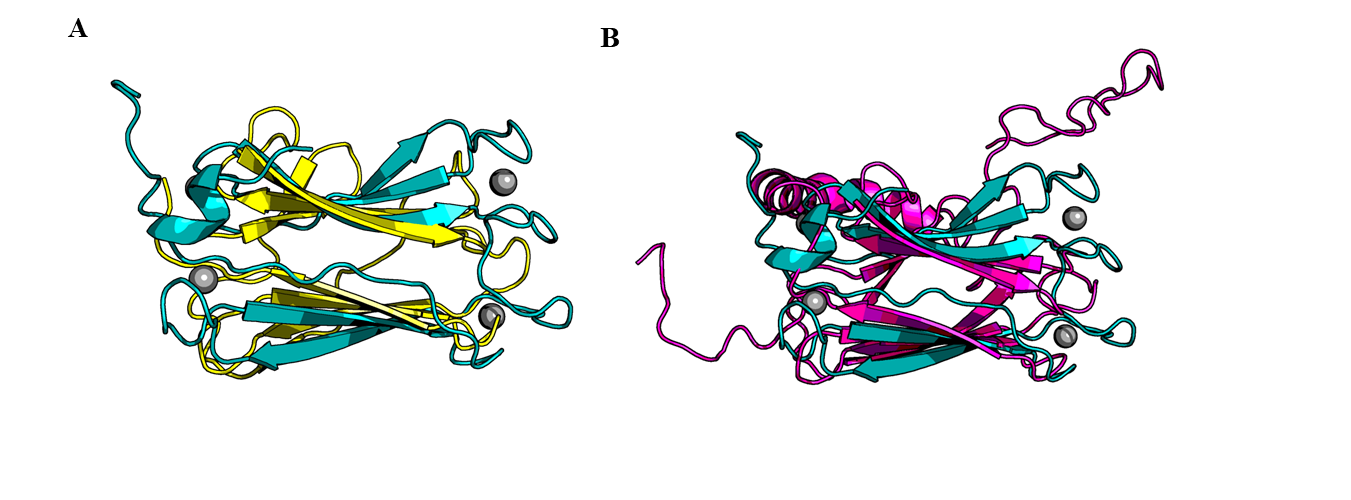

**Figure S5 AvrSr27 shares structural similarity HSP70 and DnaK .** Structural superimposition of the AvrSr27 structure (in teal, with Zn ions shown in grey), and (A) the substrate binding domains of DnaK (1dg4, yellow), and (B) HSP70 (4r5g, pink – residues 1-189 only shown). The root-mean squared deviation was 3.4 and 3.3, respectively across ~70 aligned residues.

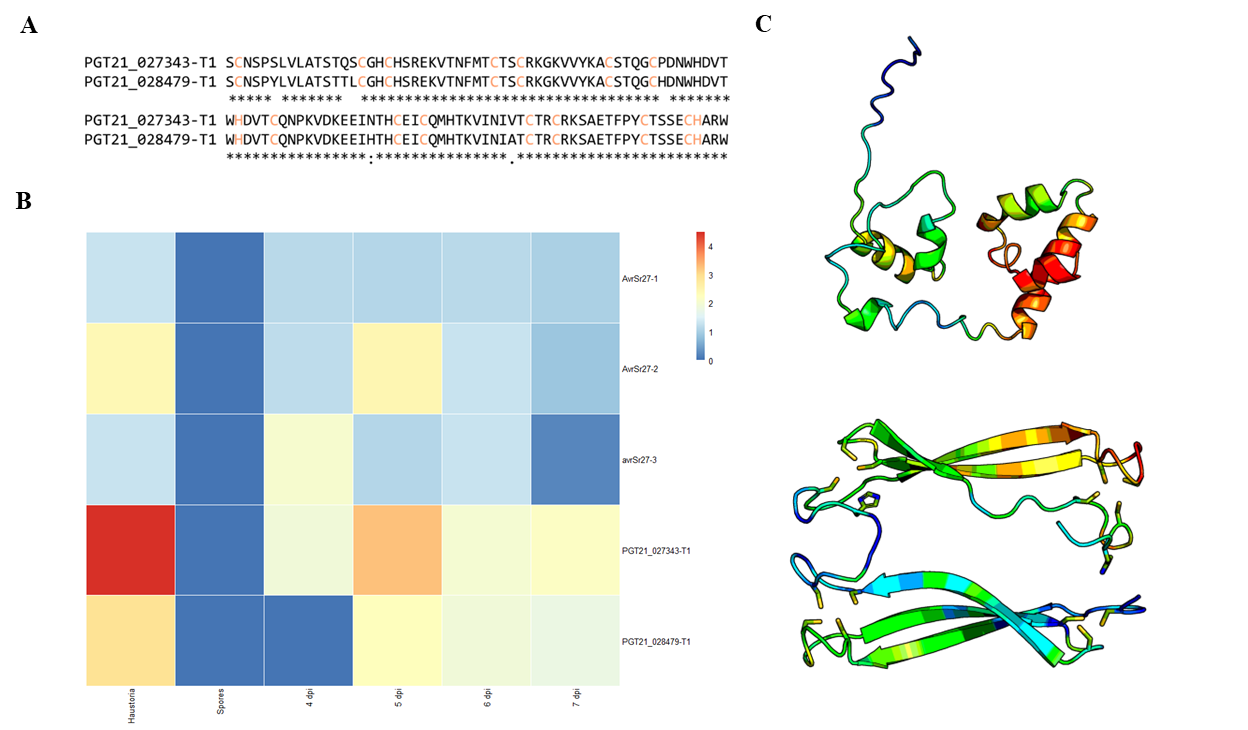

**Figure S6 The two structural homologues of AvrSr27 are expressed during infection. (A)** Multiple sequence alignment of the three allelic variants of AvrSr27 generated using MUSCLE sequence align (https://www.ebi.ac.uk/Tools/msa/muscle/). Cysteine and histidine residues conserved in AvrSr27 are highlighted in orange. **(B)** Expression levels of the two AvrSr27 structural homologues (Pgt21-027343 and Pgt21-028479) and the three *AvrSr27* variants in haustoria and germinated spores of Pgt21-0 and in infected plants at 5, 6 and 7 d post infection (dpi)(Chen et al., 2017). **(C)** AlphaFold predicted models of Pgt21-027343 (left) and Pgt21-28479 (right) shown as cartoon representations. The AlphaFold models are coloured according to pLDDT (confidence) score, where red is most confident and blue is lease confident.

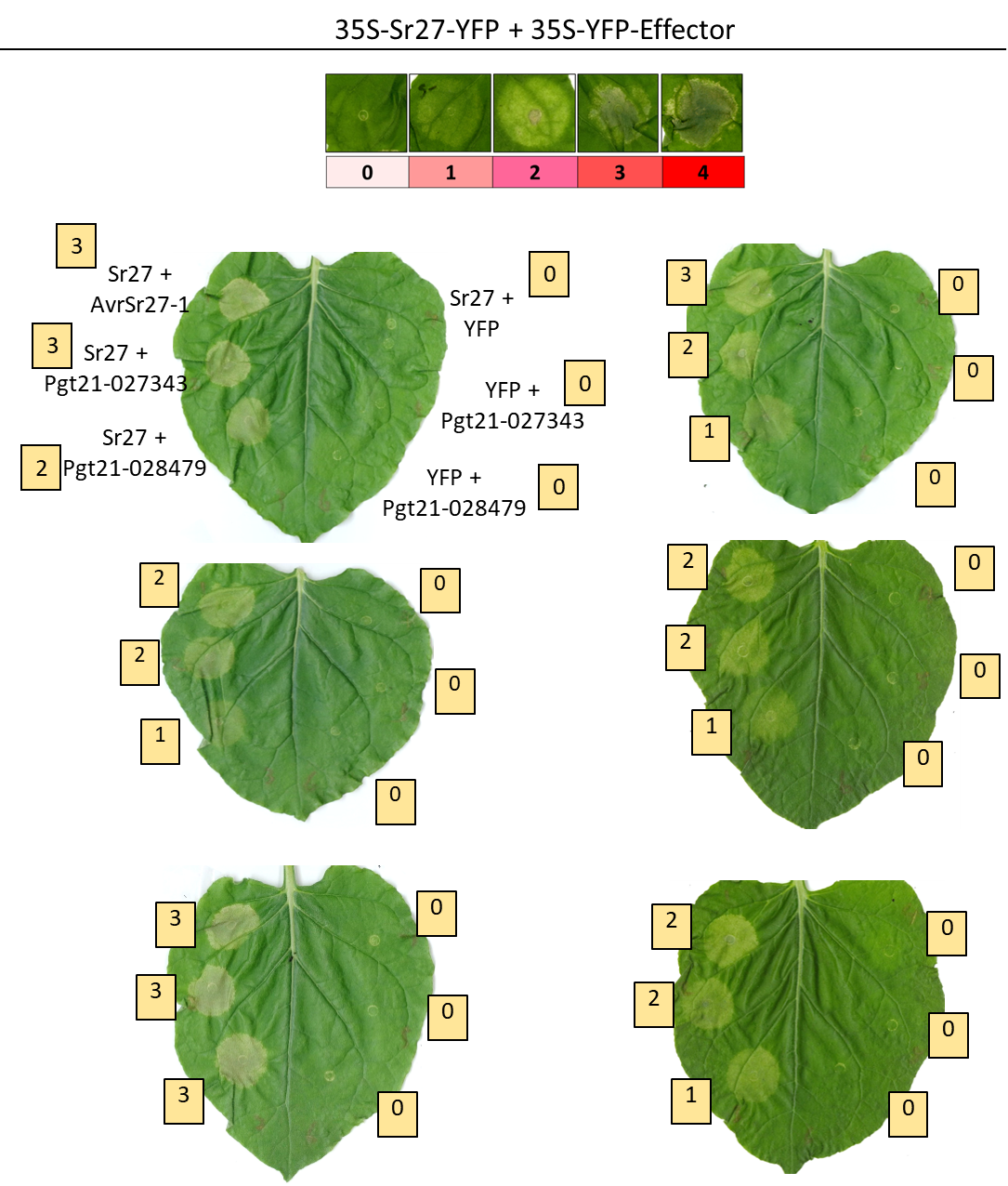

**Figure S7: Cell death response in *N. benthamiana* and *N. tabaccum* leaves transiently co-expressing Pgt21-027343 and Pgt21-028479 (AvrSr27 structural homologues). (A)** Cell death was scores according a cell death scale ranging from 0-4 (as depicted) in *N. benthamiana*. AvrSr27-1 was used as a positive control for a recognised effector-receptor interaction. Sr27, Pgt21-027343 and Pgt21-028479 alone were included as a control. The cell death scores assigned to each response correspond to the numbers shown.

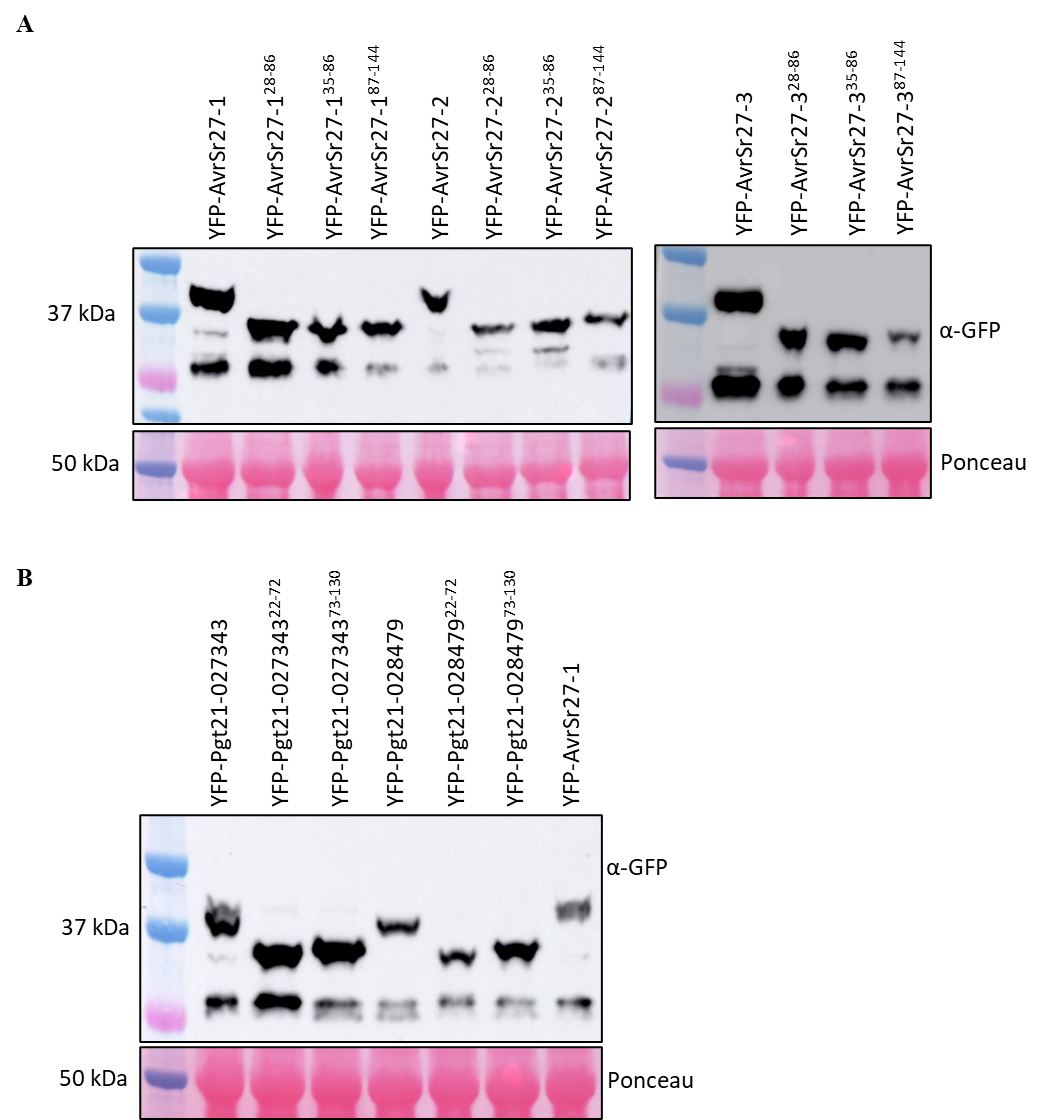

**S8: Western blot showing protein expression in *Nicotiana benthamiana* expressing the different constructs.** Protein expression was detected for (A) the three variants of AvrSr27 and the N-term (28-86 or 35-86) and C-term (87-144) fragments, and (B) the two structural homologues of AvrSr27, Pgt21-027343 and Pgt21-028479 and corresponding N- (22-72) and C-term (73-130) fragments, as derived from analysis of the AvrSr21-1 crystal structure, using anti-GFP antibodies. All constructs have an N-terminal YFP tag. Total protein was visualised by Ponceau staining method.

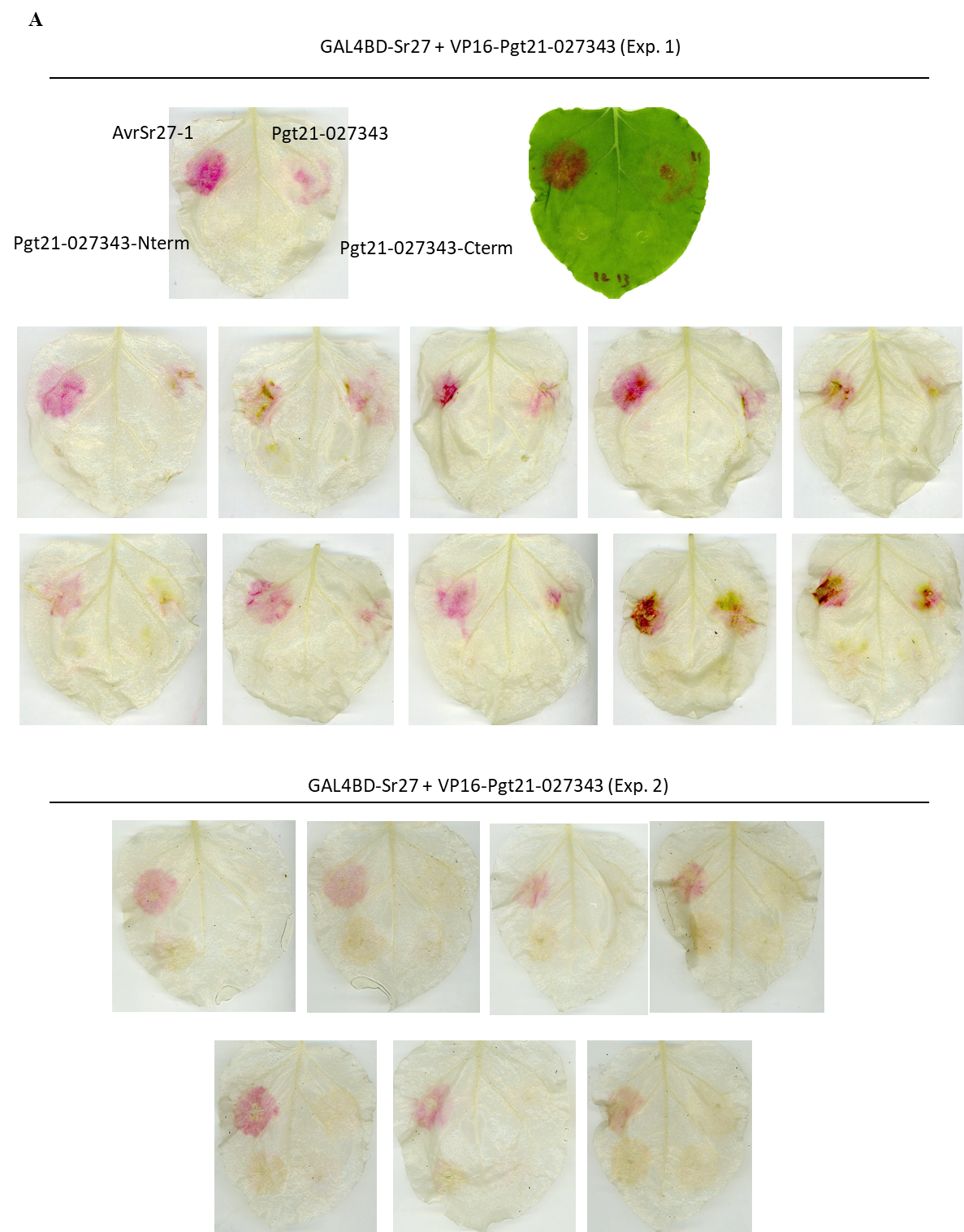

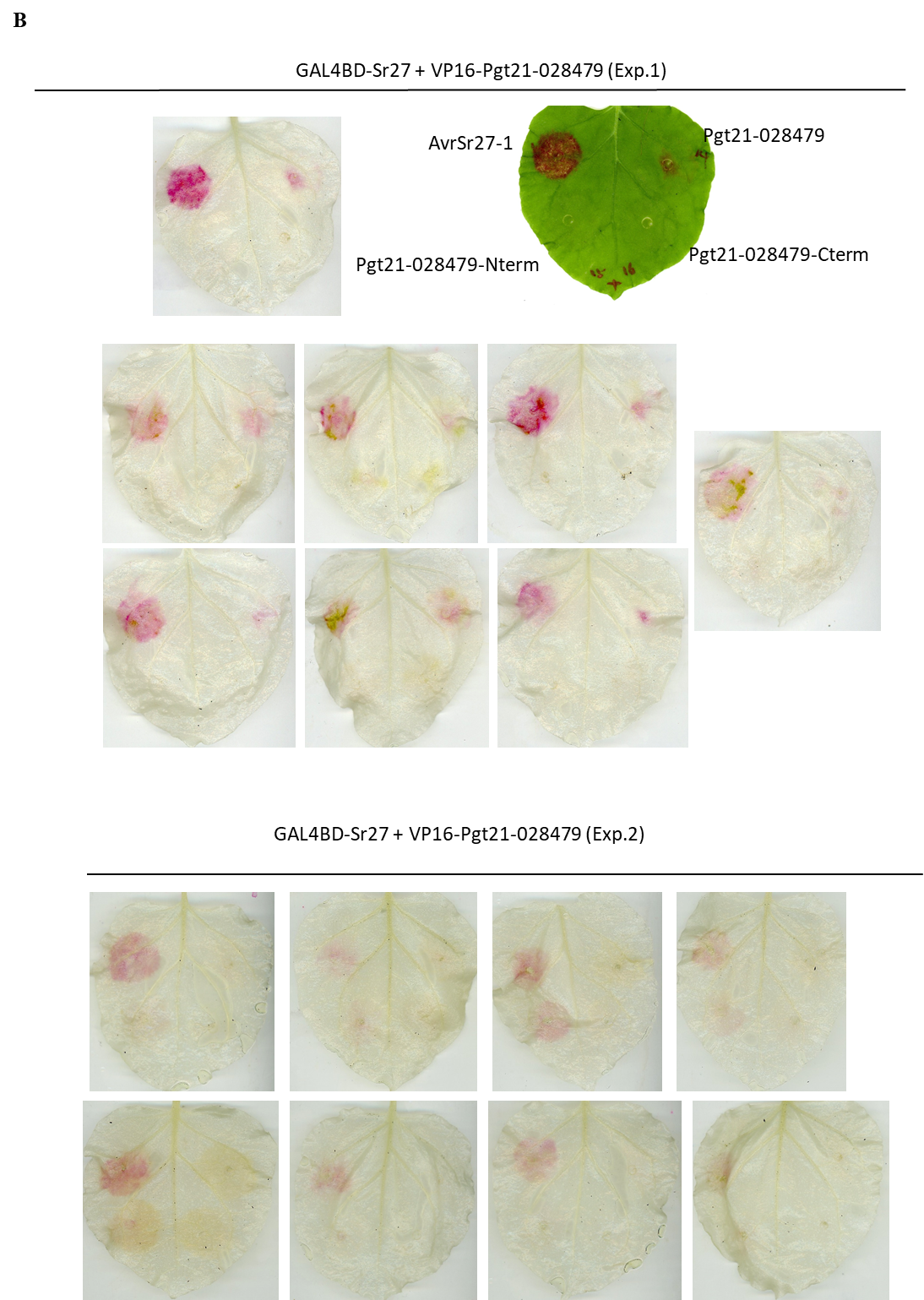

**Figure S8: *N. benthamiana* leaves transiently co-expressing the AvrSr27-like proteins and their N- and C- terminal domain splits with 35S-*RUBY* (A)** Pgt21-027343 **(B)** Pgt21-028479.

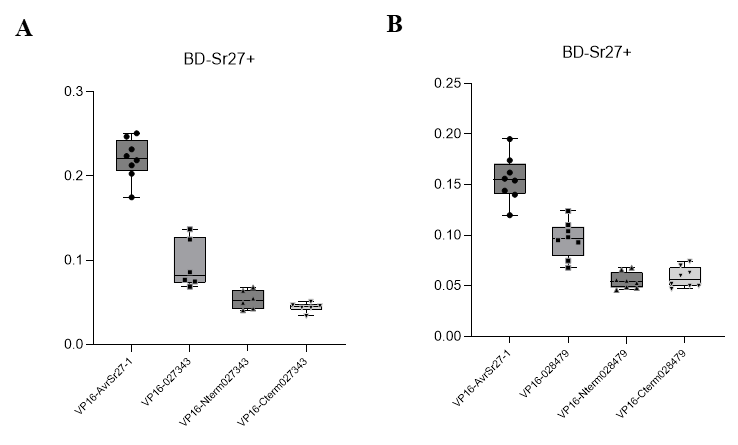

**Figure S9: Quantification of betalain (RUBY) production from *N. benthamiana* leaves transiently co-expressing the AvrSr27-like proteins and their N- and C- terminal domain splits.** Leaves as shown in Figure S8. For quantification, leaves were cleared with ethanol and leaf discs taken from infiltration sites and incubated in water for 16 hours. Absorbance at 538 nm was then measured. Common letters above columns indicate no significant difference between samples (p>0.05; one way ANOVA with posthoc Tukey HSD)

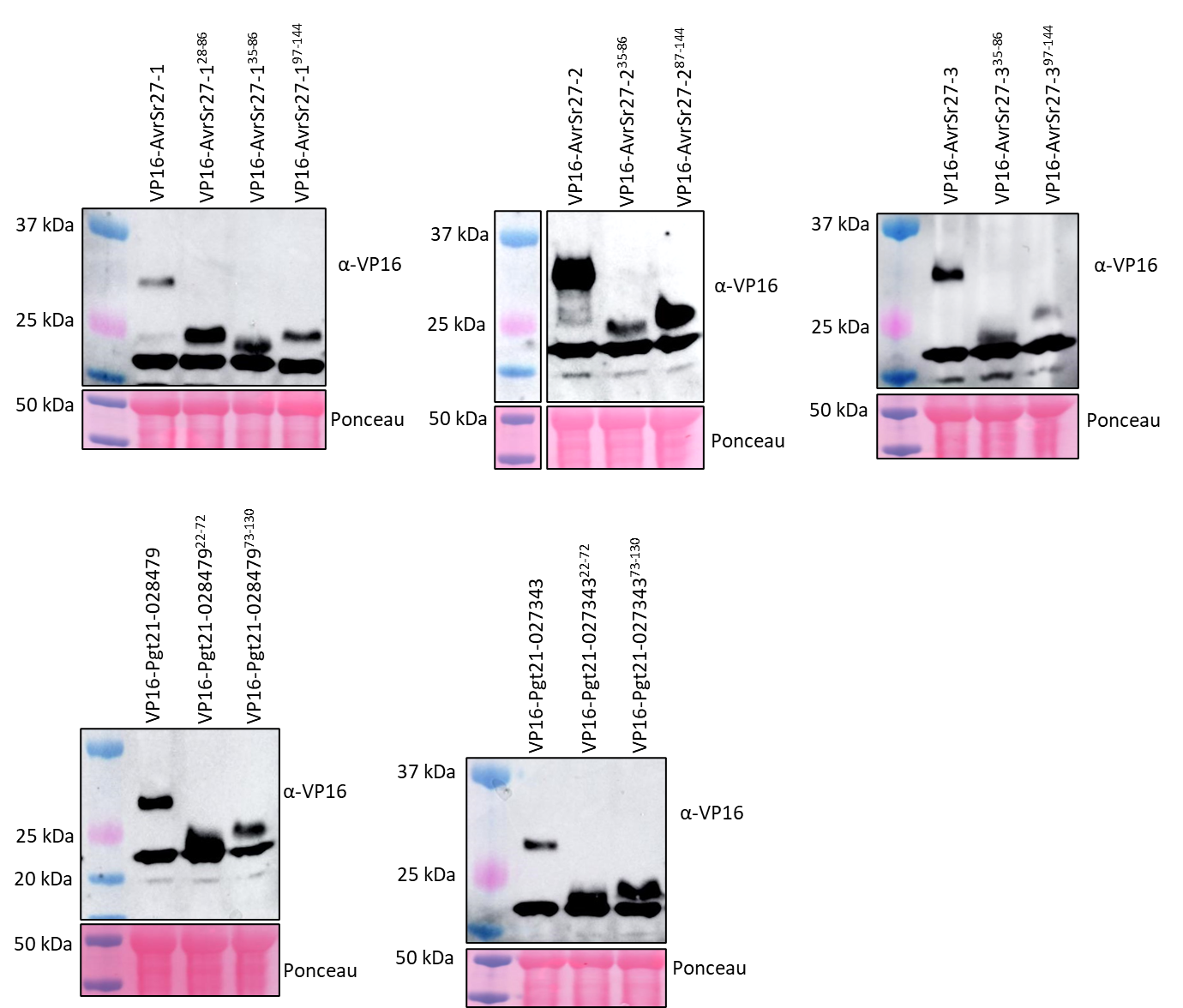

**Figure S10:** **Western blot showing protein expression in *Nicotiana benthamiana* for *in planta* two-hybrid interaction studies.** Protein expression was detected for the three variants of AvrSr27 and the N-term and C-term domains (residue boundaries are shown), and the two structural homologues of AvrSr27, Pgt21-027343 and Pgt21-028479 and corresponding N- and C-term domains, as derived from analysis of the AvrSr21-1 crystal structure. All constructs have an N-terminal VP16-tag. Total protein was visualised by Ponceau staining method.

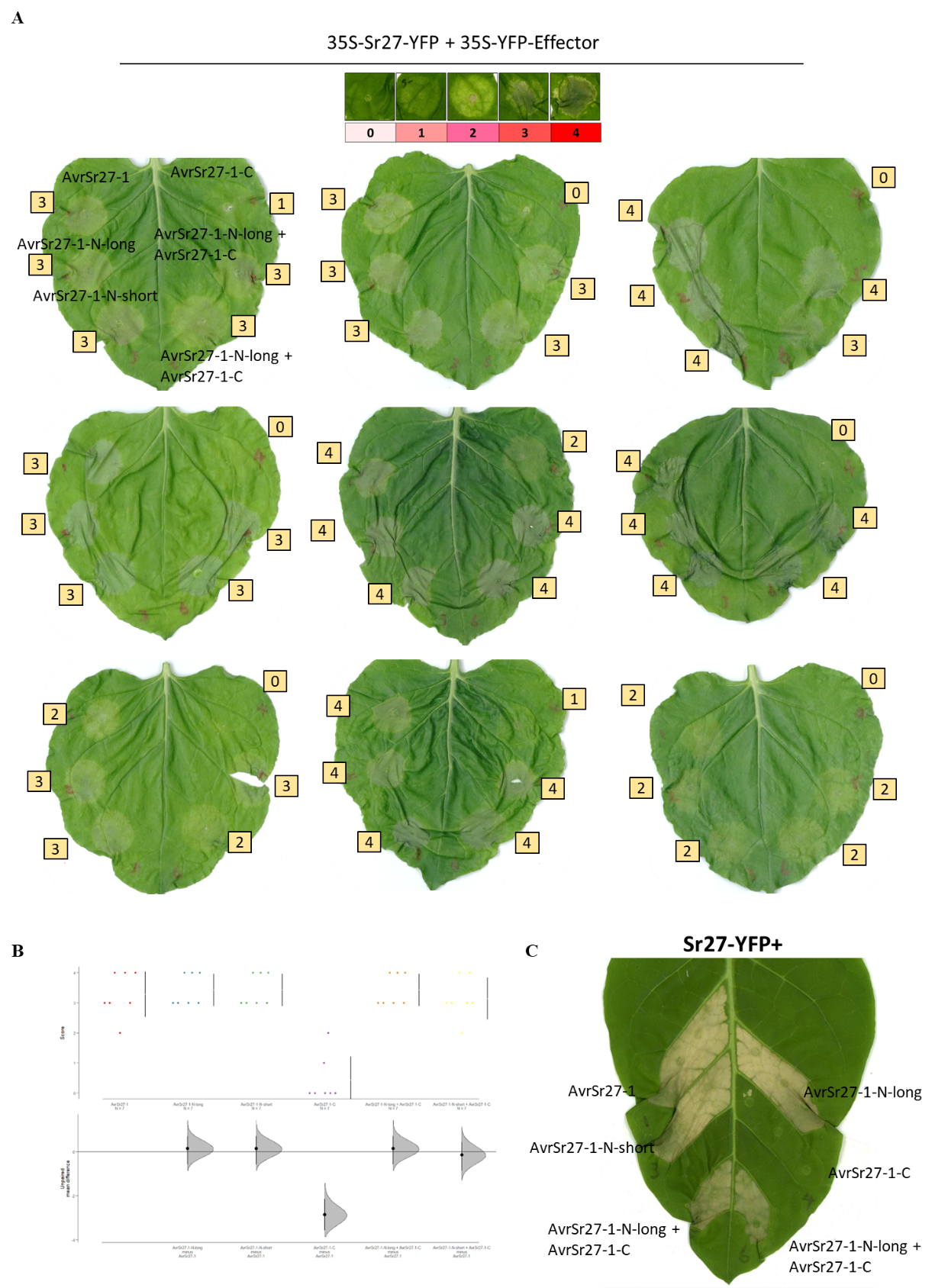

**Figure S11: Cell death response in *N. benthamiana* leaves transiently co-expressing AvrSr27-1 and the N- and C- terminal domain splits. (A)** Cell death was scored according a cell death scale ranging from 0-4 (as depicted). Full length AvrSr27-1 was used as a positive control for a recognised effector leading to cell death. The cell death scores assigned to each response correspond to the numbers shown. N-term corresponds to residues 28-86 or 35-86 and C-term residues 87-144 fragments. **(B)** Cumming estimation plot showing cell death phenotypes as represented in A with transient co-expression of the AvrSr27-1 N- and C-terminal domain splits with 35S-Sr27 in *N. benthamiana*. AvrSr27-1 was used as a positive control. The upper panels show the distribution of the observed scores for each protein where each dot represents the cell death score of an independent infiltration assay and the number of replicates per sample are indicated on the x-axis. The lower panel represents the mean differences for comparisons of variants against a shared control (AvrSr27-1 + Sr27). **(C)** Cell death response of constructs used in A transiently co-expressed with Sr27 in *Nicotiana tabacum*.

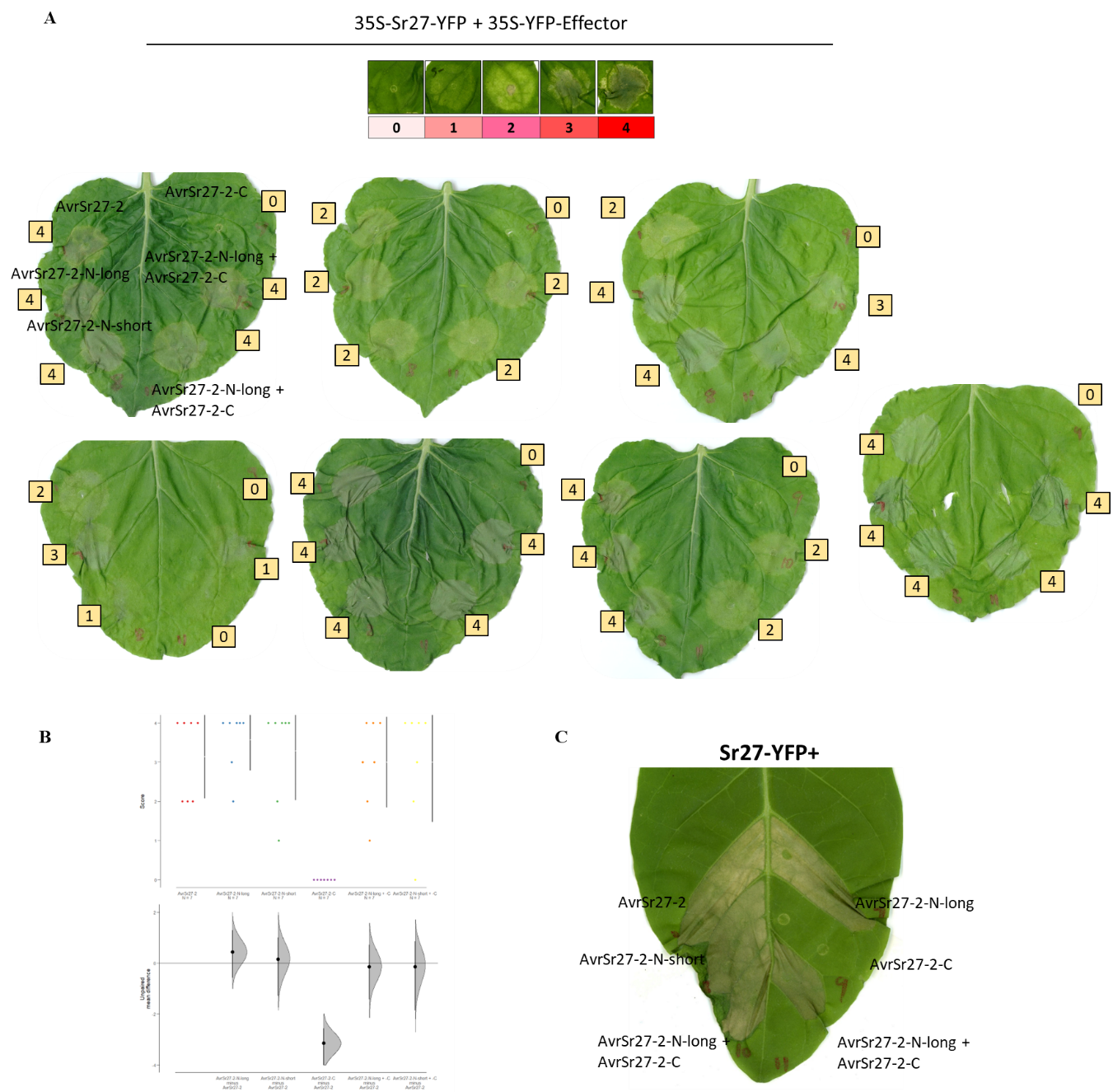

**Figure S12: Cell death response in *N. benthamiana* leaves transiently co-expressing AvrSr27-2 and the N- and C- terminal domain splits. (A)** Cell death was scored according to a cell death scale ranging from 0-4 (as depicted). Full length AvrSr27-2 was used as a positive control for a recognised effector leading to cell death. The cell death scores assigned to each response correspond to the numbers shown. N-term corresponds to residues 28-86 or 35-86 and C-term residues 87-144 fragments. **(B)** Cumming estimation plot showing cell death phenotypes as represented in A with transient co-expression of the AvrSr27-2 N- and C-terminal domain splits with 35S-Sr27 in *N. benthamiana*. AvrSr27-2 was used as a positive control. The upper panels show the distribution of the observed scores for each protein where each dot represents the cell death score of an independent infiltration assay and the number of replicates per sample are indicated on the x-axis. The lower panel represents the mean differences for comparisons of variants against a shared control (AvrSr27-2 + Sr27). **(C)** Cell death response of constructs used in A transiently co-expressed with Sr27 in Nicotiana tabacum.

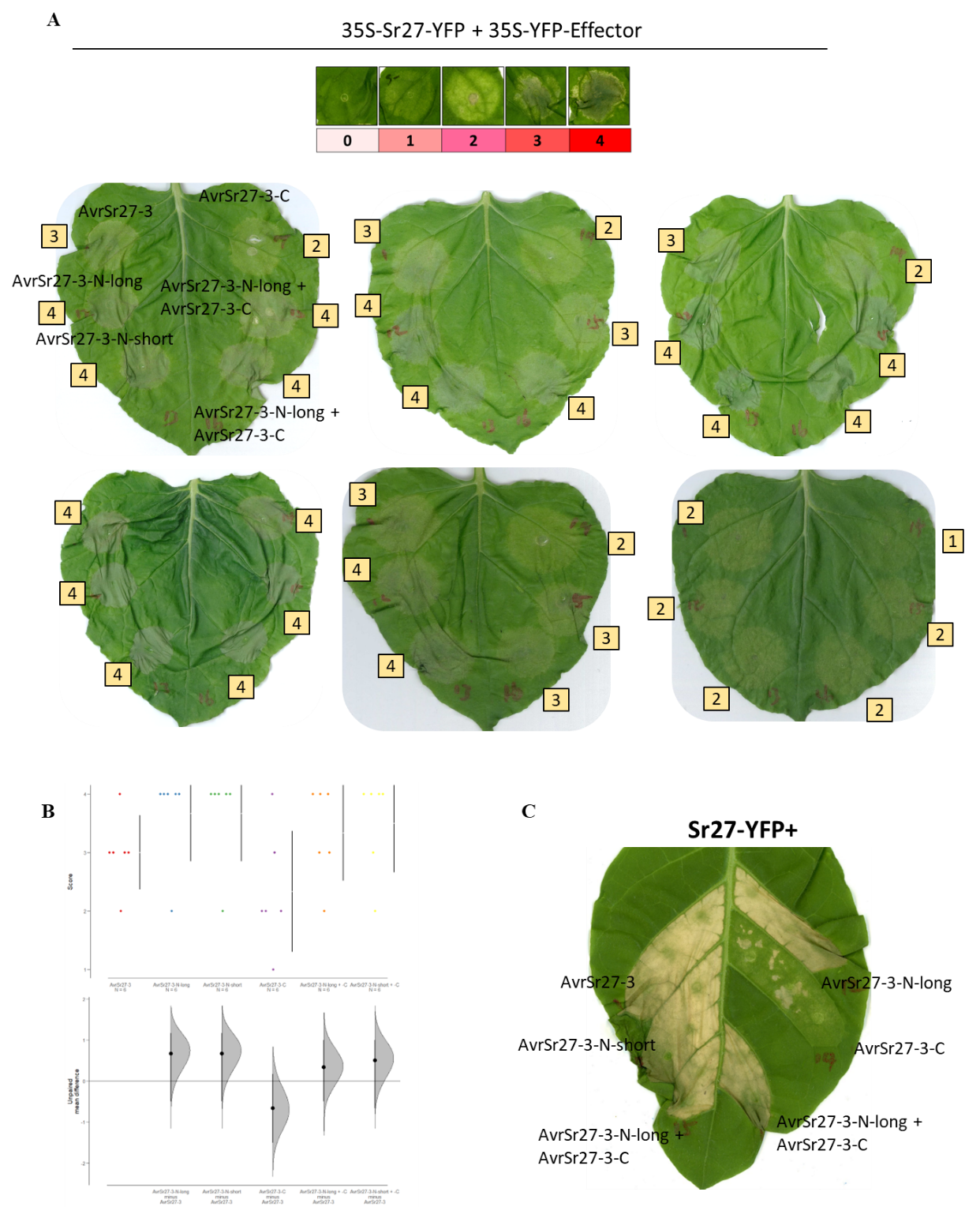

**Figure S13: Cell death response in *N. benthamiana* leaves transiently co-expressing AvrSr27-3 and the N- and C- terminal domain splits. (A)** Cell death was scored according to a cell death scale ranging from 0-4 (as depicted). Full length AvrSr27-3 was used as a positive control for a recognised effector leading to cell death. The cell death scores assigned to each response correspond to the numbers shown. N-term corresponds to residues 28-86 or 35-86 and C-term residues 87-144 fragments. **(B)** Cumming estimation plot showing cell death phenotypes as represented in A with transient co-expression of the AvrSr27-3 N- and C-terminal domain splits with 35S-Sr27 in *N. benthamiana*. AvrSr27-3 was used as a positive control. The upper panels show the distribution of the observed scores for each protein where each dot represents the cell death score of an independent infiltration assay and the number of replicates per sample are indicated on the x-axis. The lower panel represents the mean differences for comparisons of variants against a shared control (AvrSr27-3 + Sr27). **(C)** Cell death response of constructs used in A transiently co-expressed with Sr27 in *Nicotiana tabacum*.

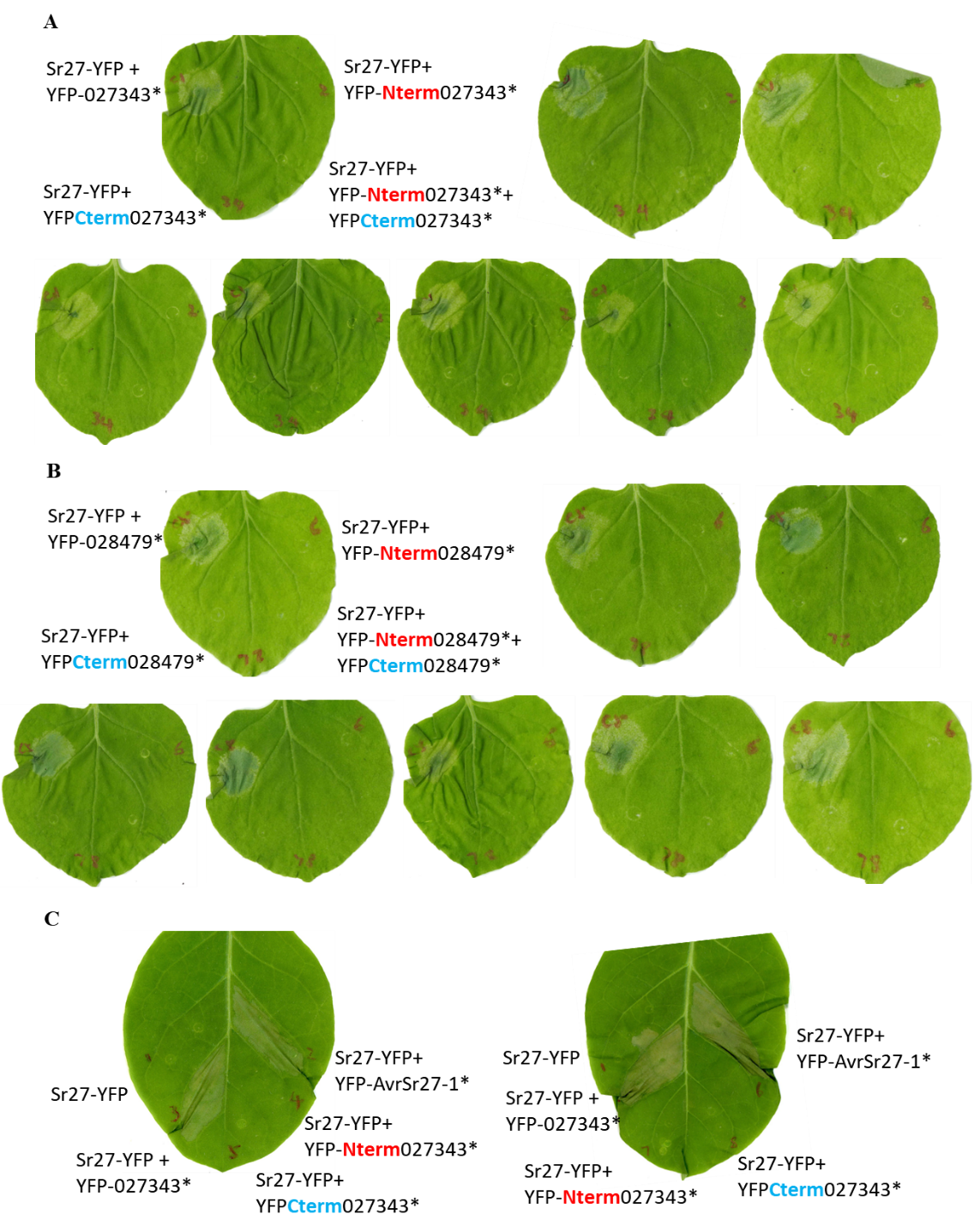

**Figure S14: Cell death response in *N. benthamiana* leaves transiently co-expressing Pgt21-027343 and Pgt21-028479 N- and C- terminal domain splits with Sr27. (A, B)** Full length Pgt21-027343 and Pgt21-028479 were used as a positive control for cell death. N-term corresponds to residues 22-72 or 73-130 and C-term residues 87-144 fragments. **(C)** Cell death response of constructs used in A and B transiently co-expressed with Sr27 in *Nicotiana tabacum*.

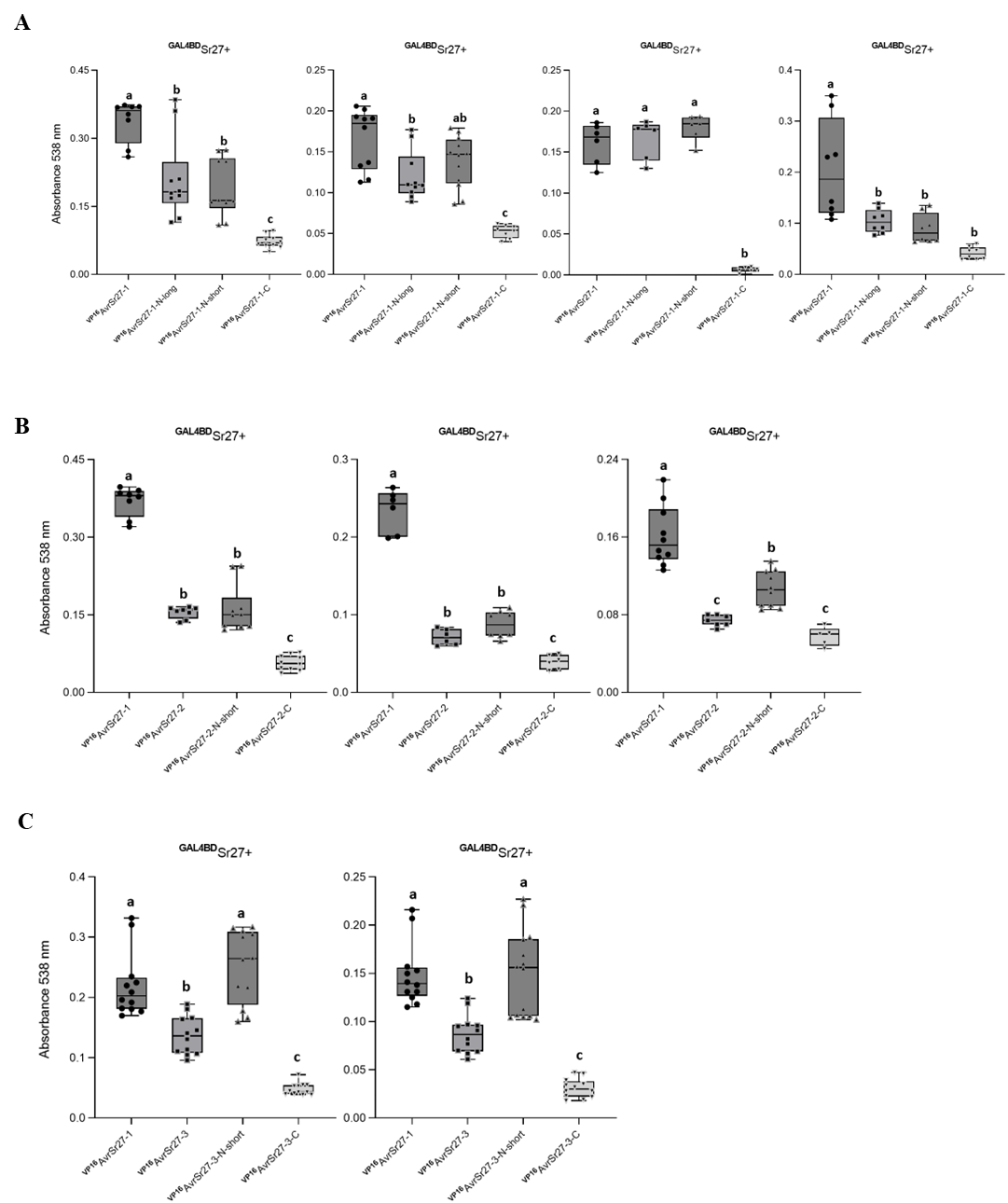

**Figure S15: The N-terminus of the AvrSr27 variants directly interacts with Sr27.** Quantification of betalain (RUBY) production from each infiltration site was achieved by clearing leaves with ethanol and incubating leaf disks from infiltration sites in water for 16 hours. Absorbance at 538 nm was then measured. Common letters above columns indicate no significant difference between samples (p>0.05; one way ANOVA with posthoc Tukey HSD). All leaves used for quantification are shown in Fig S9A-C.

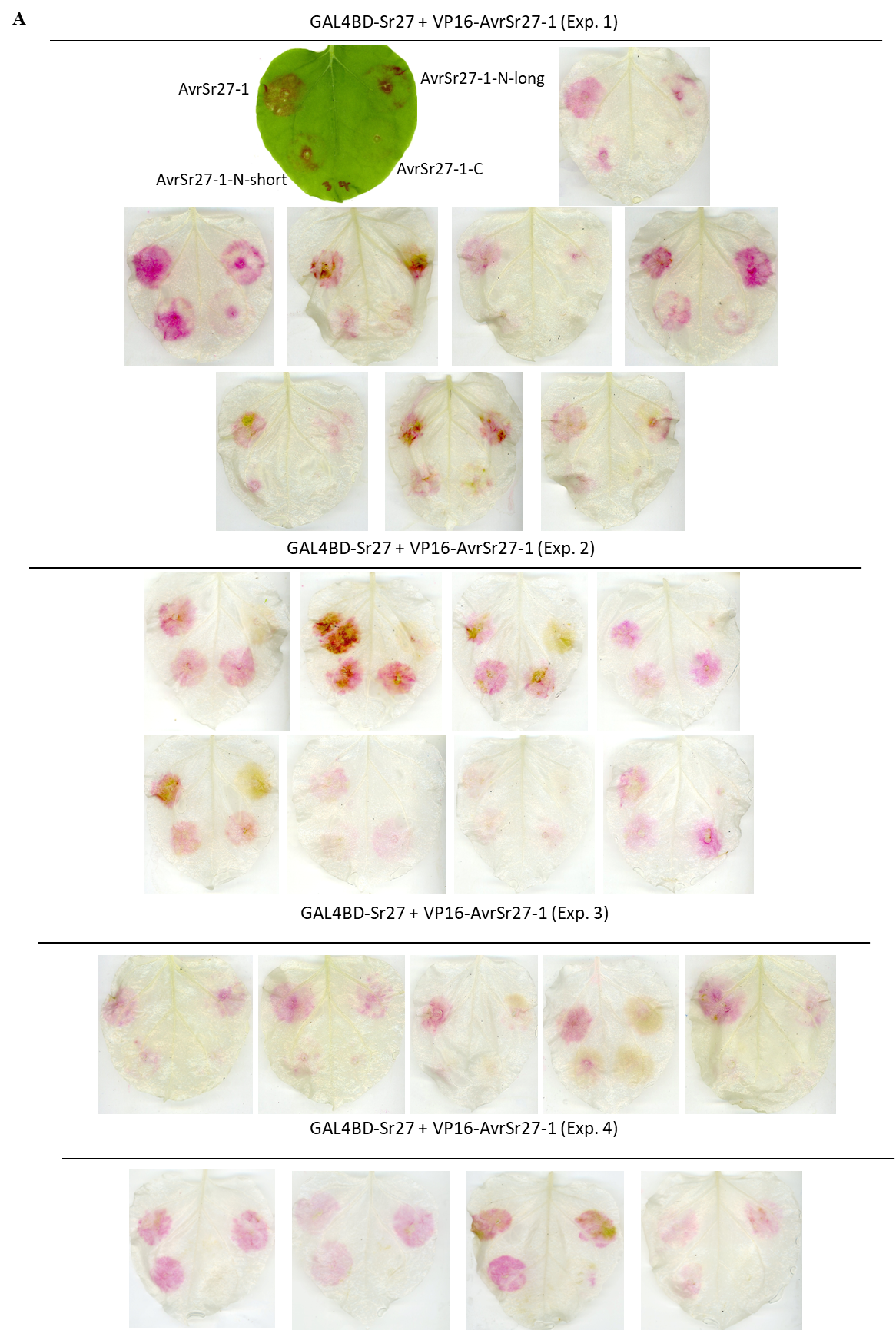

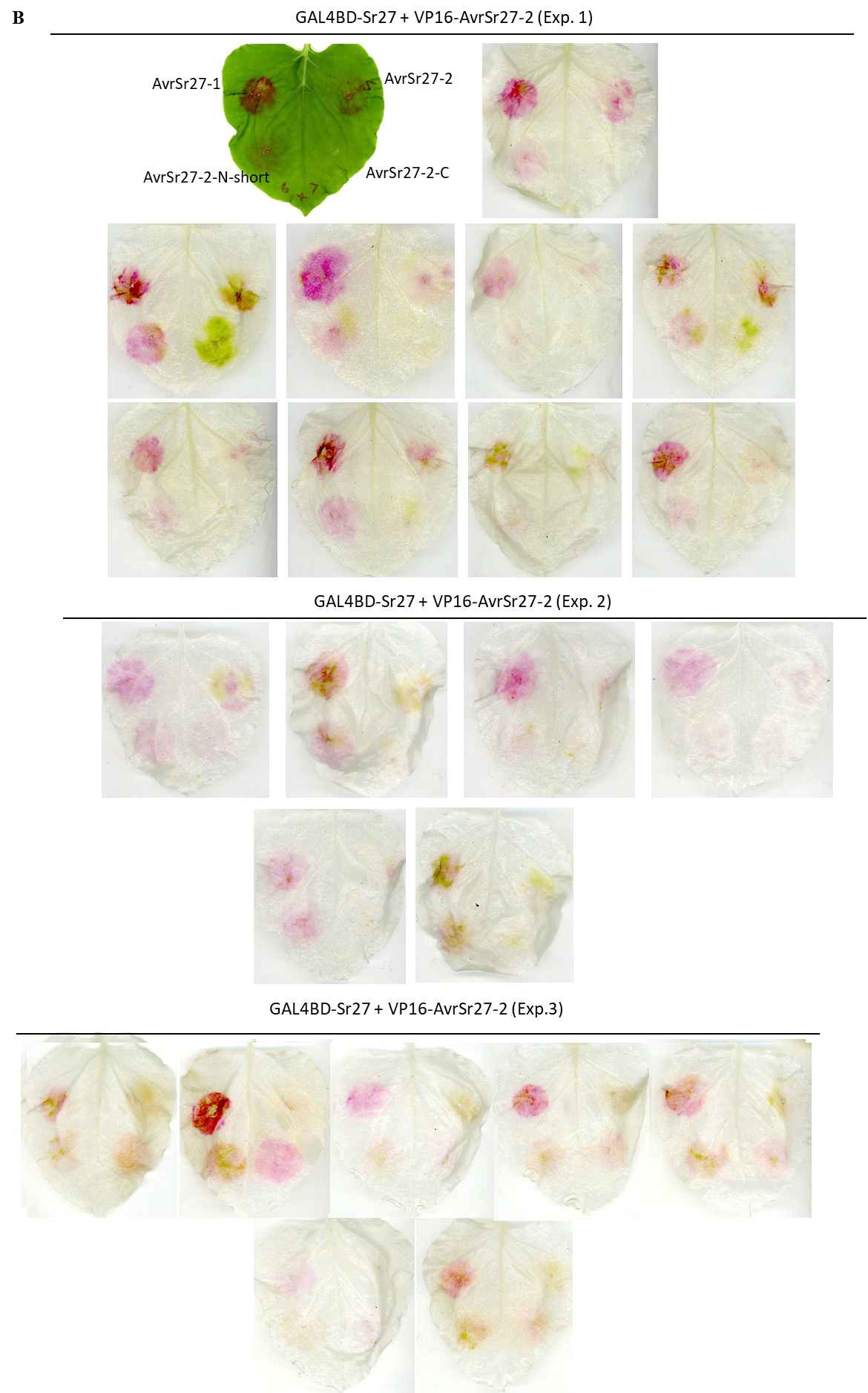

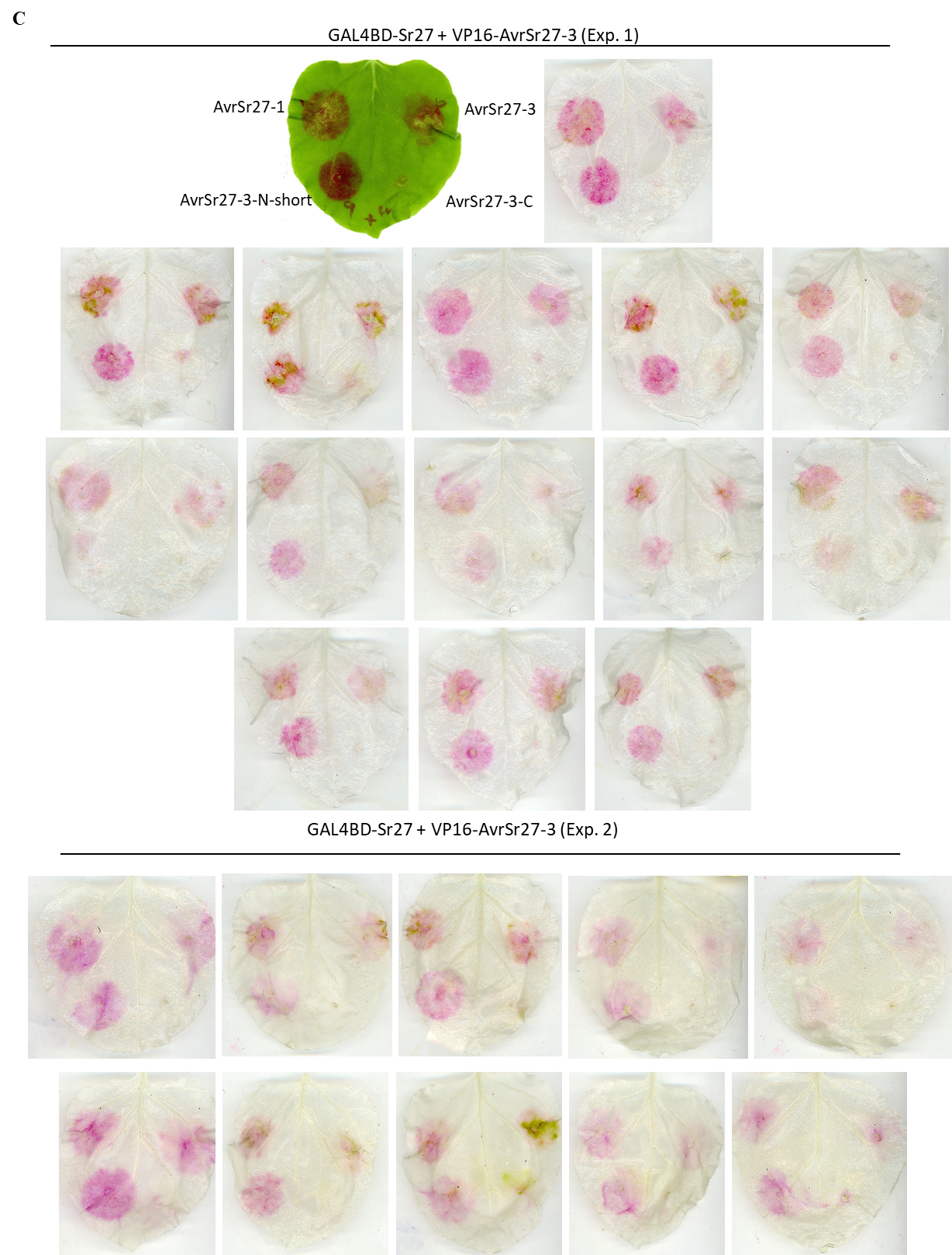

**Figure S16: *N. benthamiana* leaves transiently co-expressing AvrSr27 variants and the N- and C- terminal domain splits with *RUBY*, as showed in Fig S8. (A)** AvrSr27-1 **(B)** AvrSr27-2, and **(C)** AvrSr27-3.
